# Spatially Constrained Monte Carlo Permutation Test Reveals Diffusion Changes Near Stress Granules

**DOI:** 10.64898/2026.09.17.752460

**Authors:** Elizaveta Korunova, Vitali Sikirzhytski, Jeffery L. Twiss, Michael Shtutman, Paula Vasquez

## Abstract

Intracellular diffusion is inherently heterogeneous, yet single-particle tracking (SPT) analyses are often summarized using cell-wide average parameters that can obscure localized effects. Here, we tracked 40-nm genetically encoded multimeric (GEM) nanoparticles during stress granule (SG) formation and developed SPaCe-MC (Spatially Constrained Monte Carlo permutation test), a statistical framework that generates cytoplasm-specific null models to test whether diffusion associated with a specific cellular structure differs from that expected in the surrounding heterogeneous cytoplasm. Across three SG-inducing conditions, including oxidative stress, DDX3 inhibition, and combined treatment, bulk cytoplasmic analyses revealed distinct responses ranging from increased nanoparticle mobility to increased subdiffusive behavior. In contrast, SPaCe-MC consistently detected a local diffusion constraint in SG-associated regions relative to their treatment-matched cytoplasmic background, revealing a conserved local diffusion effect despite divergent global cytoplasmic responses. Together, our findings establish SPaCe-MC as a framework for identifying compartment-specific diffusion changes in heterogeneous cellular environments.

## Introduction

The physical properties of the cytoplasm are evolutionarily optimized and are tightly integrated with fundamental cellular processes, including protein synthesis and degradation^1^, cell cycle progression^2^, and cytoskeletal remodeling^3^. Consistent with this functional role, single-particle tracking (SPT) studies have shown that cytoplasmic diffusion and macromolecular crowding are dynamically regulated during nutrient deprivation^4^, cellular senescence^5^, or under compressive stress^6^. However, diffusion is inherently heterogeneous across cellular environments^7–10^, and ensemble- and time-averaged mean squared displacement (MSD) analyses, which are commonly used to characterize diffusion parameters, typically yield cell-wide average diffusion parameters that may mask biologically relevant heterogeneity^11,12^. To characterize spatial variation in diffusion, recent studies have employed approaches ranging from kernel regression with statistical bootstrapping to reconstruct diffusion maps in giant fungal cells^8^ to Bayesian inference–based methods, including DiffMAP-GP^13^, InferenceMAP^14^, and TRamWAy^15^. These approaches infer spatially resolved diffusion landscapes from SPT tracks and provide powerful frameworks for visualizing and quantifying where diffusion varies within the cytoplasm. However, this raises a complementary biological question: does diffusion within a specific cytoplasmic structure differ significantly from that expected in the surrounding heterogeneous environment?

Liquid–liquid phase separation (LLPS) provides a biologically relevant setting for investigating this question. By compartmentalizing proteins and/or nucleic acids into dynamic condensates ranging from micron-sized assemblies to extended network-like structures^16^, LLPS generates spatially distinct intracellular environments that are expected to differentially regulate molecular diffusion^17^. Stress granules (SGs), one of the most extensively studied RNA–protein condensates, assemble in the cytoplasm during cellular stress to reorganize translation, signaling, and RNA metabolism^18^. Consistent with their dynamic organization, SGs are commonly described as heterogeneous condensates composed of less mobile cores surrounded by more dynamic shell regions, with their inner dynamics regulated by ATP-dependent RNA helicases^19^.

Measurements using genetically encoded multimeric nanoparticles (GEMs, 40 nm) and μNS particles (50–150 nm) have shown that stress-induced SG formation accompanied by polysome disassembly increases cytoplasmic diffusivity across multiple length scales^20^. However, these measurements primarily describe cell-scale responses and do not directly address whether diffusion differs between SG-associated regions and the surrounding cytoplasm. Probing condensate properties *in vivo* remains challenging, although GEMs have been proposed as a tool that could be optimized for this purpose^17^. While SGs have been shown to reduce the accessible space for ∼29-nm quantum dots^21^, local changes in the diffusion of inert fluorescent probes near SGs have not yet been characterized using SPT. More broadly, determining whether observed differences between SG-associated and surrounding cytoplasmic regions reflect genuine compartment-specific effects requires accounting for the limited number of tracks within small cytoplasmic regions, finite track lengths, uneven spatial sampling, and the stochastic nature of diffusion^11,12^.

Here, we introduce SPaCe-MC, a spatially constrained Monte Carlo permutation framework designed to test whether diffusion associated with a predefined cellular structure differs from that expected within the surrounding heterogeneous cytoplasm. SPaCe-MC adapts Monte Carlo randomization, widely used to assess statistical significance in microscopy colocalization analyses^22^, to spatially resolved diffusion measured by SPT. Rather than constructing spatial diffusion maps, SPaCe-MC tests a compartment-level hypothesis by comparing diffusion parameters estimated from tracks associated with the original region of interest with a null distribution generated from randomly sampled cytoplasmic regions. This approach enables population-level testing of region-specific diffusion changes while accounting for stochastic particle motion and spatial sampling variability.

We applied SPaCe-MC to GEM tracks during SG formation to determine whether SG-associated diffusion changes persist across conditions that produce distinct bulk cytoplasmic responses. We examined NaAsO₂-induced oxidative stress, RK-33-mediated inhibition of the ATP-dependent DEAD-box helicase DDX3, a key regulator of SG remodeling^23^, and combined RK-33/NaAsO₂ treatment. These conditions induce SG formation through distinct cellular perturbations and, in the case of RK-33, produce condensates with altered RNA composition^18^ as well as molecular dynamics^23^. By comparing SG-associated diffusion with the treatment-specific cytoplasmic background, these conditions allowed us to test whether a common SG-associated diffusion phenotype could be distinguished from inducer-dependent changes in bulk cytoplasmic dynamics.

Using SPaCe-MC in combination with bulk cytoplasmic diffusion analysis, simulations, and super-resolution imaging, we show that distinct SG-inducing conditions produce divergent changes in bulk cytoplasmic diffusion, yet SG-associated regions (defined here as local subcellular regions spatially defined by SGs, rather than strictly as the SG interior) consistently exhibit constrained GEM mobility. These SG-associated regions occupy only ∼2–4% of the segmented cytoplasmic area, and super-resolution imaging shows that most GEM nanoparticles classified as SG-associated reside within ∼100 nm of SG boundaries, indicating that the observed diffusion changes occur primarily near the condensate periphery. Together, these findings demonstrate how SPaCe-MC can resolve region-specific diffusion changes that are obscured by bulk cytoplasmic measurements and reveal that SGs generate a conserved local physical environment despite inducer-dependent remodeling of the surrounding cytoplasm.

## Results

### Global cytoplasmic diffusion analysis reveals inducer-dependent cytoplasmic responses

First, we estimated the overall effect of SG formation on cytoplasmic nanoparticle dynamics in U2OS cells subjected to three SG-inducing conditions: oxidative stress induced by sodium arsenite (NaAsO₂), DDX3 inhibition using RK-33, or combined RK-33/NaAsO₂ (Combo) treatment. Cytoplasmic dynamics were monitored using 40-nm genetically encoded multimeric nanoparticles (GEMs) expressed under a tight tetracycline-inducible promoter^11^ together with the SG marker G3BP1-mCherry. Two-dimensional live-cell imaging was performed to track GEM motion and quantify MSD-based bulk cytoplasmic diffusion^4–6^ (Fig. 1A).

**Figure 1.**
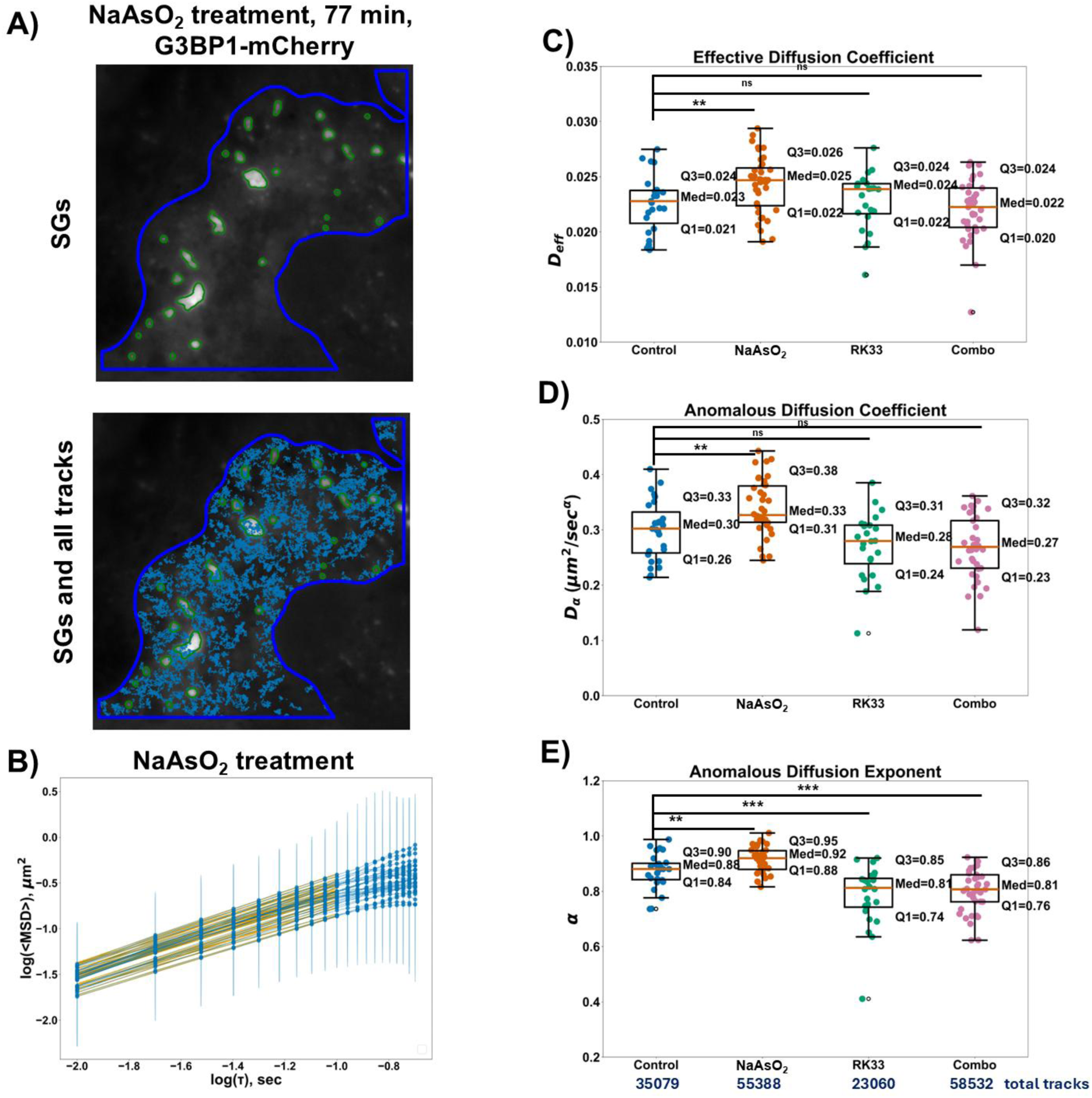
Different cytoplasmic diffusion responses at the 40-nm scale under SG inducers. **(A)** Representative U2OS cell expressing G3BP1-mCherry. Green outlines indicate SGs, and the blue outline marks the G3BP1-positive cytoplasm. Top, G3BP1 image; bottom, overlaid GEM single-particle tracking (SPT) tracks. **(B)** Time- and ensemble-averaged mean squared displacement (MSD) was estimated from all cytoplasmic tracks pooled over one video and plotted against lag time (τ) on a log–log scale for NaAsO₂ treatment. Blue, experimental data; yellow, fit to the anomalous Brownian motion model, MSD = 4 Dα t^a^. **(C–E)** Diffusion parameters obtained from anomalous Brownian motion fits to the first 100 ms of the MSD curve (R² ≥ 0.9): (C) effective diffusion coefficient (Deff), (D) anomalous diffusion coefficient (Dα), and (E) anomalous exponent (α). Cells were imaged at the onset of NaAsO₂ treatment (1 h after RK-33 or DMSO treatment) and analyzed after 30 min of imaging, corresponding to robust NaAsO₂-induced SG formation. In Figs. 1C-E, each dot corresponds to the diffusion parameter estimated from all GEM tracks detected within one video (75 × 75 µm field of view).

Diffusion parameters were obtained by fitting MSD curves (Fig. 1B) to an anomalous diffusion model (MSD = 4 D_α_ t^a^), yielding the generalized diffusion coefficient (D_α_) and anomalous exponent (α), where α = 1 corresponds to Brownian diffusion and α < 1 to subdiffusive motion arising from confinement or molecular crowding^11^ (See Methods). We first compared our results with previously reported cytoplasmic responses to NaAsO₂ treatment measured using GEMs^20,21^. Consistent with stress-induced cytoplasmic fluidization^20^, NaAsO₂-treated cells exhibit higher values of D_eff_, D_α_, and α than untreated controls (Fig. 1C–E). In addition, apparent diffusion coefficients (D) estimated under the assumption of Brownian diffusion (α = 1) were also higher in NaAsO₂-treated cells than in controls (Fig. S1A–B; see Methods)^4,20^. In contrast, paired imaging of the same cell culture before and after NaAsO₂ treatment yielded the opposite trend, with decreases in D_eff_, D_α_, and α following treatment (Fig. S2). This finding is consistent with a previous paired-cell study^21^ but contrasts with both our independently imaged populations (Fig. 1C–E; Fig. S1) and the previous report of stress-induced cytoplasmic fluidization^20^. Because repeated imaging may contribute to this discrepancy, all subsequent analyses were performed using independently imaged cell populations.

We next examined whether pre-treatment with RK-33 affected the changes in cytoplasmic dynamics during SG formation. We found that RK-33 treatment alone or pre-treatment before arsenite did not increase either D_α_ or D_eff_ relative to untreated cells (Fig. 1C–E). Analysis of anomalous diffusion parameters further revealed a decrease in the anomalous exponent α (∼0.9 → ∼0.8), indicating a subtle increase in GEM subdiffusion under conditions involving RK-33 treatment (Fig. 1D).

Consistent with recent findings in the field, these observations demonstrate that SG formation can be accompanied by different changes in cell-wide cytoplasmic diffusion. Polysome disassembly and cytoplasmic mRNA levels have been shown to determine the diffusional response to stress^20,24^, while polysome disassembly together with SG formation has been proposed as a prerequisite for cytoplasmic fluidization in mammalian cells under NaAsO_2_ treatment^20^. However, because bulk cytoplasm diffusion measurements can reflect the combined effects of SG-associated processes, including polysome collapse^20^ and redistribution of RNA between condensates and the cytoplasm^18^, as well as broader stress-dependent cellular responses^25^, they cannot determine whether diffusion is specifically altered within SG-associated regions.

### SPaCe-MC framework design and statistical validation

To determine whether SG-associated regions exhibit diffusion properties distinct from the surrounding cytoplasm, we developed SPaCe-MC (Spatially Constrained Monte Carlo permutation test), a region-based statistical framework in which SGs serve as regions of interest (ROI) (Fig. 2A; see Methods). SPaCe-MC enables the detection of region-specific diffusion changes while improving the precision of diffusion-parameter estimates through the pooling of tracks across cells. For each treatment, diffusion parameters estimated from tracks associated with the original SG masks are compared with a null distribution generated by repeatedly relocating those masks within the surrounding cytoplasm and pooling the sampled tracks across cells.

**Figure 2.**
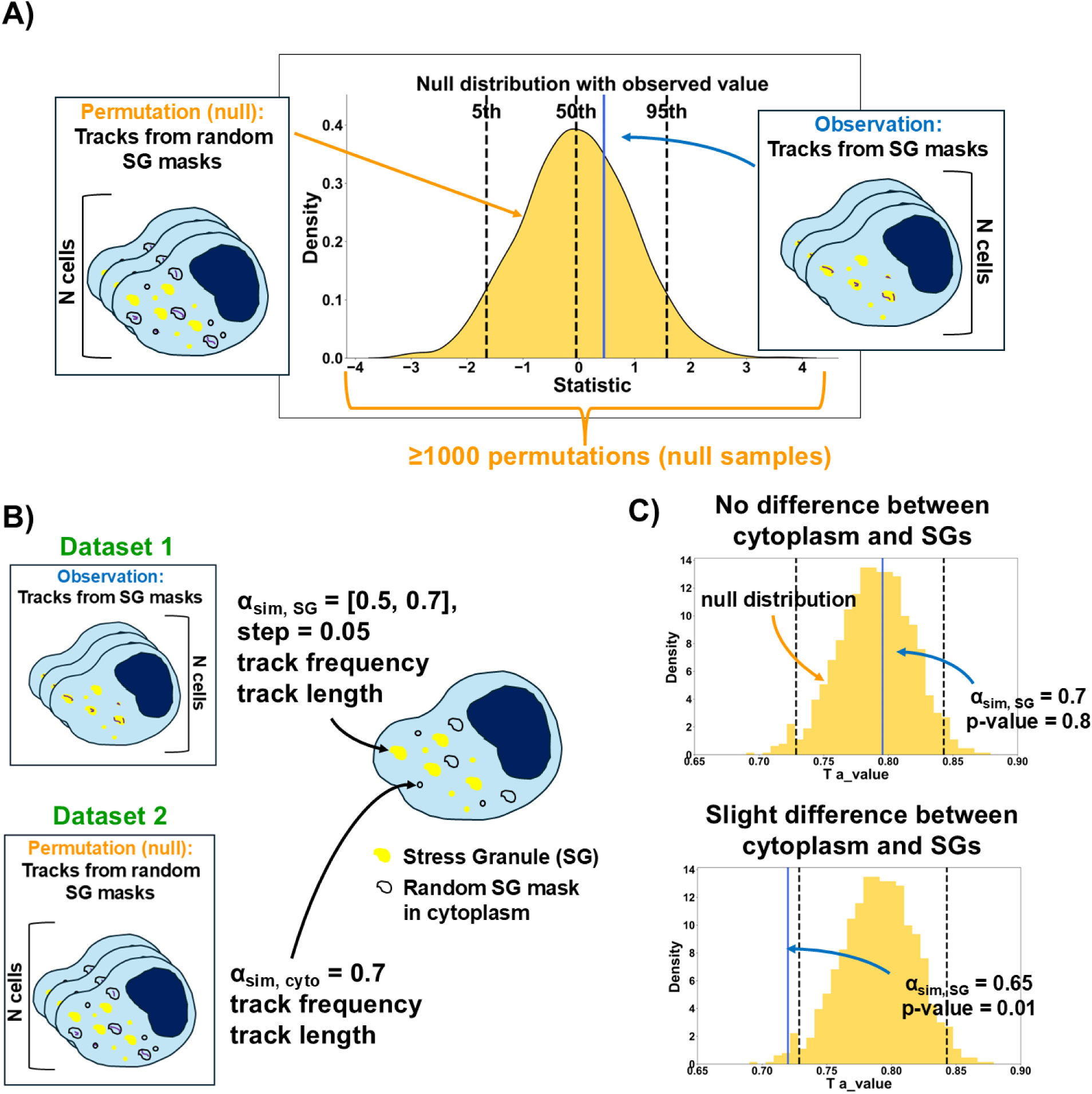
SPaCe-MC workflow for detecting ROI-associated local changes in diffusion relative to the surrounding cytoplasm. **(A)** Schematic of the SPaCe-MC workflow. A null distribution is generated from ≥1000 permutations by randomly relocating SG masks throughout the cytoplasm while excluding existing SG regions across videos. For each permutation, SG masks are randomized within each segmented cytoplasmic region, and the resulting tracks are pooled across videos to estimate a single diffusion parameter. **(B)** Simulated dataset used for method validation. fBm tracks reproduced the experimentally observed track numbers and durations. Cytoplasmic tracks were simulated with H = 0.35 (α = 0.70), whereas SG-associated tracks were simulated with H = 0.25–0.35 (α = 0.50–0.70). **(C)** Representative null distribution generated from simulated fBm tracks. The histogram shows diffusion parameters obtained from 1000 cytoplasmic permutations. Black lines indicate the 2.5th and 97.5th percentiles, and the blue line indicates the diffusion parameter measured within the simulated SG region.

To validate SPaCe-MC, we quantified the sampling characteristics of SG-associated tracks and tracks sampled using permuted cytoplasmic masks across videos within each treatment. We then estimated the expected performance of SPaCe-MC using simulated track datasets with sampling characteristics matched to the experimental data. Through simulations, we distinguished between parameters used to generate simulated tracks and parameters estimated from track data. In the fractional Brownian motion (fBM) simulations, the Hurst parameter (H) and input diffusion coefficient (D_i_) defined the simulated motion. The resulting tracks, as well as experimental tracks, were analyzed using MSD-based models to estimate the anomalous exponent (α), anomalous diffusion coefficient (D_α_), effective diffusion coefficient (D_eff_), and, where indicated, the apparent Brownian diffusion coefficient (D). Results of simulations were used to evaluate the statistical power and false-positive rate (FPR) of the permutation test (Fig. 3; Figs. S4–S15; see Methods for details).

**Figure 3.**
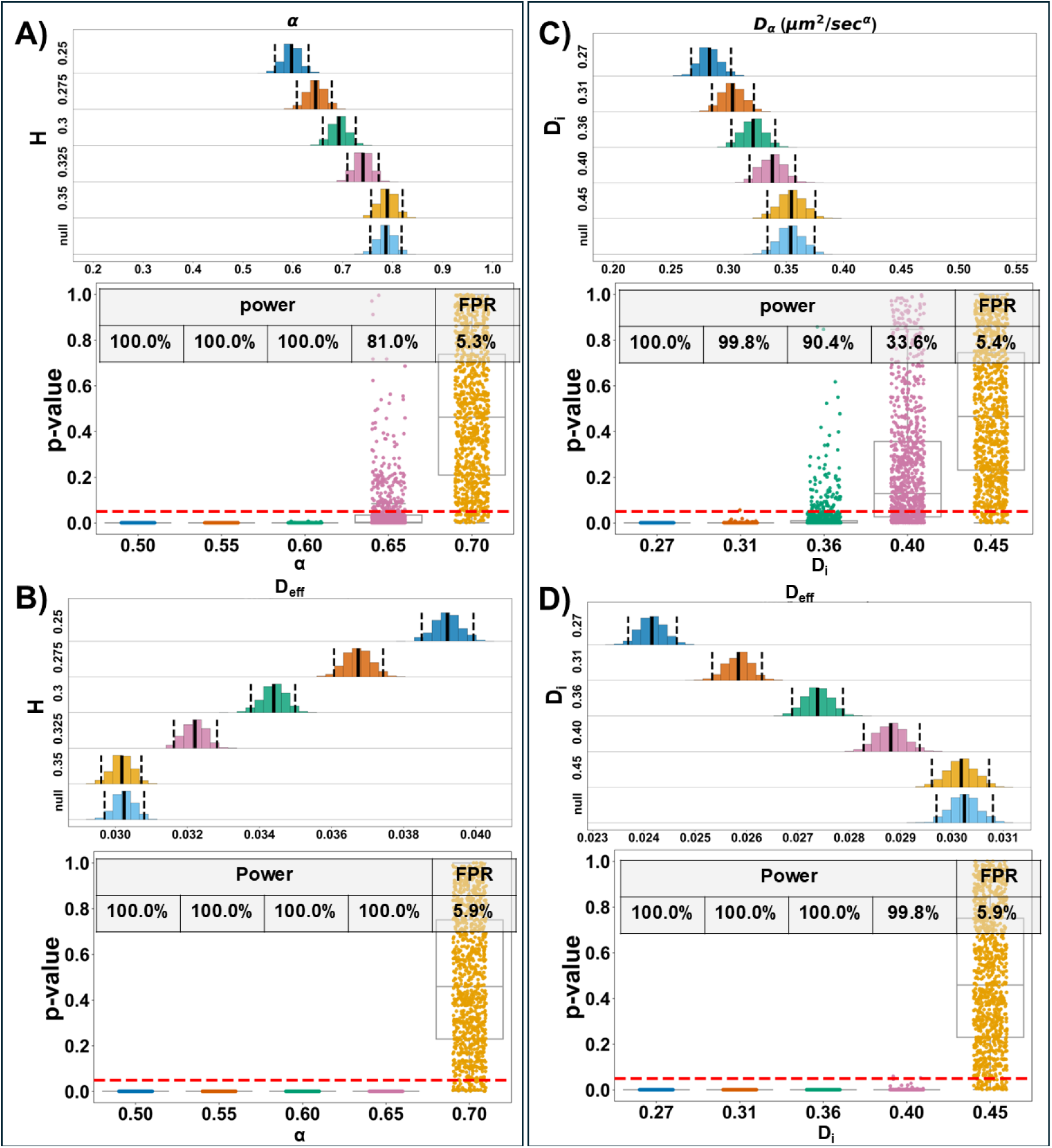
False-positive rate and power of SPaCe-MC using simulated datasets matched to experimental track statistics (Combo treatment). **(A–B)** Joyplots of the estimated **(A)** α and **(B)** Deff together with boxplots of permutation-test p values. Simulations were generated by varying H = 0.25–0.35 (Y-axis; corresponding to α = 0.50–0.70) while keeping Di = 0.45 μm²/s constant. **(C–D)** Joyplots of the estimated **(C)** Dα and **(D)** Deff together with boxplots of permutation-test p values. Simulations were generated by varying Di = 0.27–0.45 μm²/s (Y-axis) while keeping H = 0.35 constant. **(A–D)** Simulated datasets reproduced the experimental distributions of track number and duration for the Combo treatment (Fig. S3) but contained no spatial organization. Null datasets were generated with H = 0.35 (α = 0.70) and Di = 0.45 μm²/s, whereas observation datasets varied either H or Di. Statistical power and FPR were estimated from 1,000 simulated datasets (37 trajectory groups per dataset; p < 0.05).

We found that the FPR for estimates of α and Dα was approximately 5–6% at sample sizes matching the Combo treatment (Fig. 3A-C). Statistical power decreased as the simulated parameters approached the null, reaching 81% for Δα = 0.05 and 90.4% for ΔDi = 0.09 μm²/s.

We also evaluated how permutation-test performance depends on sample size and matching between the observed and null distributions. We considered two sampling schemes: matched sampling, in which the observed and null datasets had matched track-number and track-duration distributions, and experimental sampling, in which the observed and null datasets reproduced the SG-associated and permutation-derived sampling characteristics, respectively. The matched-sampling simulations therefore evaluated SPaCe-MC performance when sampling differences between the observed and null datasets were removed, whereas the experimental-sampling simulations evaluated performance under the sampling differences present in the experimental data. Under matched sampling, the FPR remained between 4% and 6% (Figs. S4–S9), whereas statistical power for detecting Δα = 0.05 decreased from 80–90% under NaAsO₂- and Combo-like sampling (∼1,400 and 1,000 tracks per permutation) to ∼33% under the lower-sampled RK-33 condition (∼200 tracks per permutation). Under experimental sampling, differences between the SG-associated and permutation-derived sampling distributions increased the FPR to as much as 33% for the NaAsO₂- and RK-33-like data sets (Figs. S12–S15). Interestingly, simulations also showed that both D and D_eff_ depend on α and D_α_, complicating their interpretation as purely viscosity-related parameters, although D_eff_ exhibited greater statistical power than α and D_α_ (Fig. 3B–D; Figs. S4–S15).

We next generated simulated null distributions to characterize the effects of sampling and diffusion parameters on permutation-test performance (Figs. S16–S18). We systematically varied track number, track duration, input diffusion coefficient, and Hurst parameter. As expected, null-distribution width depended strongly on the number of tracks per permutation, with smaller samples producing greater uncertainty in diffusion parameter estimates (Fig. S18). At ∼20 tracks per permutation, corresponding to sampling within an SG mask in one video, this uncertainty substantially limited meaningful comparisons at the individual-cell level.

Together, these results support the use of SPaCe-MC for population-level comparisons under sufficiently sampled, matched-null conditions. Statistical power increased with track number and effect size, whereas mismatched sampling increased the FPR, emphasizing the importance of appropriate sampling and null-model construction for robust statistical inference.

### SPaCe-MC reveals localized diffusion changes associated with stress granules

We then applied SPaCe-MC to characterize diffusion changes associated with SG regions (Fig. 4). Across all SG-inducing treatments, GEMs in SG-associated regions showed consistently lower α, D_α_, and D_eff_ than expected from the surrounding cytoplasm (Δα ≈ 0.05–0.1, p ≈ 0.01– 0.04; ΔD_α_ ≈ 0.04 μm²/s^α, p ≈ 0.002–0.04; Fig. 4A-C). Because the validation simulations showed that differences between SG-associated and permutation-derived sampling characteristics can inflate the FPR, we interpreted the experimental p values in the context of the treatment-specific simulation results. The strength of statistical support, however, depended on treatment-specific sampling characteristics. For the NaAsO₂ sampling scheme, a more stringent threshold of p < 0.01 for α and D_α_ yielded an FPR of 12.6%, whereas p < 0.001 for D_eff_ yielded an FPR of 2.9%. Track frequency within SG regions did not differ significantly from the null distribution, except under NaAsO₂ treatment, where GEM tracks were significantly depleted from SG regions (Fig. 4D).

**Figure 4.**
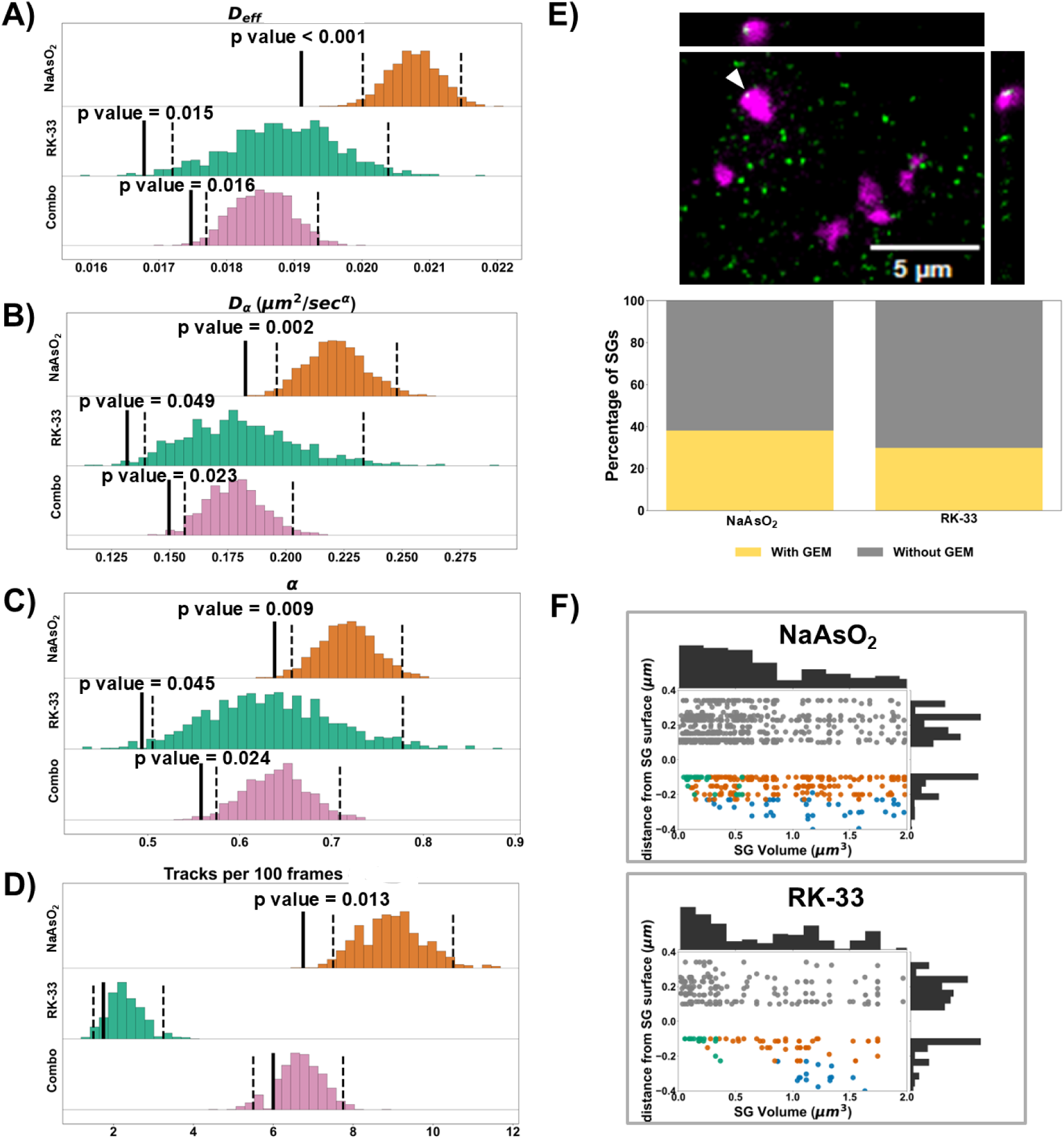
SG-associated local diffusion and GEM localization at stress granules. **(A–D)** Null distributions of **(A)** Deff, **(B)** Dα, **(C)** α, and **(D)** track frequency generated by permutation of SG masks across segmented U2OS cells. Black lines indicate the 2.5th and 97.5th percentiles (95% interval), and the blue line the observed value measured within SG masks. The observed value was estimated from tracks pooled across videos within the original SG masks. For each permutation, tracks were pooled across cells from one permuted mask within each segmented cytoplasmic region in each video. **(E) Top:** Representative cropped z-stack of U2OS cells showing G3BP1-labeled SGs (magenta) and GEM nanoparticles (green) with orthogonal views. White arrows indicate the SG that is included in the orthogonal view. **Bottom:** Fraction of GEM nanoparticles with center-of-mass coordinates within segmented SG boundaries. **(F)** Spatial distribution of GEM nanoparticles relative to SGs. Joint plot with marginal distributions showing particles in SG interior (blue; two erosions), periphery (orange), neighboring cytoplasm (gray; three dilations), and SGs too small for subdivision (green).

Thus, despite distinct bulk cytoplasmic responses to different SG-inducing conditions, SPaCe-MC revealed a common SG-associated diffusion signature across all treatments, indicating a local reduction in GEM mobility across distinct SG-inducing conditions.

### Nanoparticle probes preferentially localize to stress granule peripheries

To determine the spatial relationship between GEM nanoparticles and SGs, we performed STED imaging of SGs labeled with anti-G3BP1 STAR RED antibodies together with confocal imaging of GEMs in cells treated with either NaAsO_2_ or RK-33 (Fig. 4E). GEM positions were estimated from particle center-of-mass coordinates and compared with segmented SG volumes. Across both conditions, approximately 30–40% of GEMs localized within SG boundaries.

To further characterize the spatial distribution of GEMs relative to SG architecture, SG volumes were subdivided into interior, peripheral, and neighboring regions. The interior was defined as the SG volume remaining after two successive erosions, the peripheral region as the SG shell excluded from the interior, and the neighboring region as the cytoplasmic area surrounding the SG (Fig. 4F). Analysis of GEM localization revealed a preference for SG peripheries over interior regions, with most GEMs residing within approximately 100 nm of the SG surface. Thus, GEM interactions with G3BP1-defined SGs appear to occur predominantly near the SG boundary, potentially involving both the peripheral SG region and the adjacent cytoplasm.

### SPaCe-MC null distributions capture treatment-dependent cytoplasmic diffusion backgrounds

Because SPaCe-MC evaluates SG-associated diffusion relative to a treatment-specific cytoplasmic background, we next examined whether the resulting null distributions capture the distinct cytoplasmic diffusion states observed across SG-inducing conditions. To facilitate comparison across treatments using a standardized ROI geometry, we generated treatment-specific null distributions from experimental data using randomly positioned masks consisting of 10 circular regions (area = 1.7 μm²) instead of SG masks (Fig. 5A–D).

**Figure 5.**
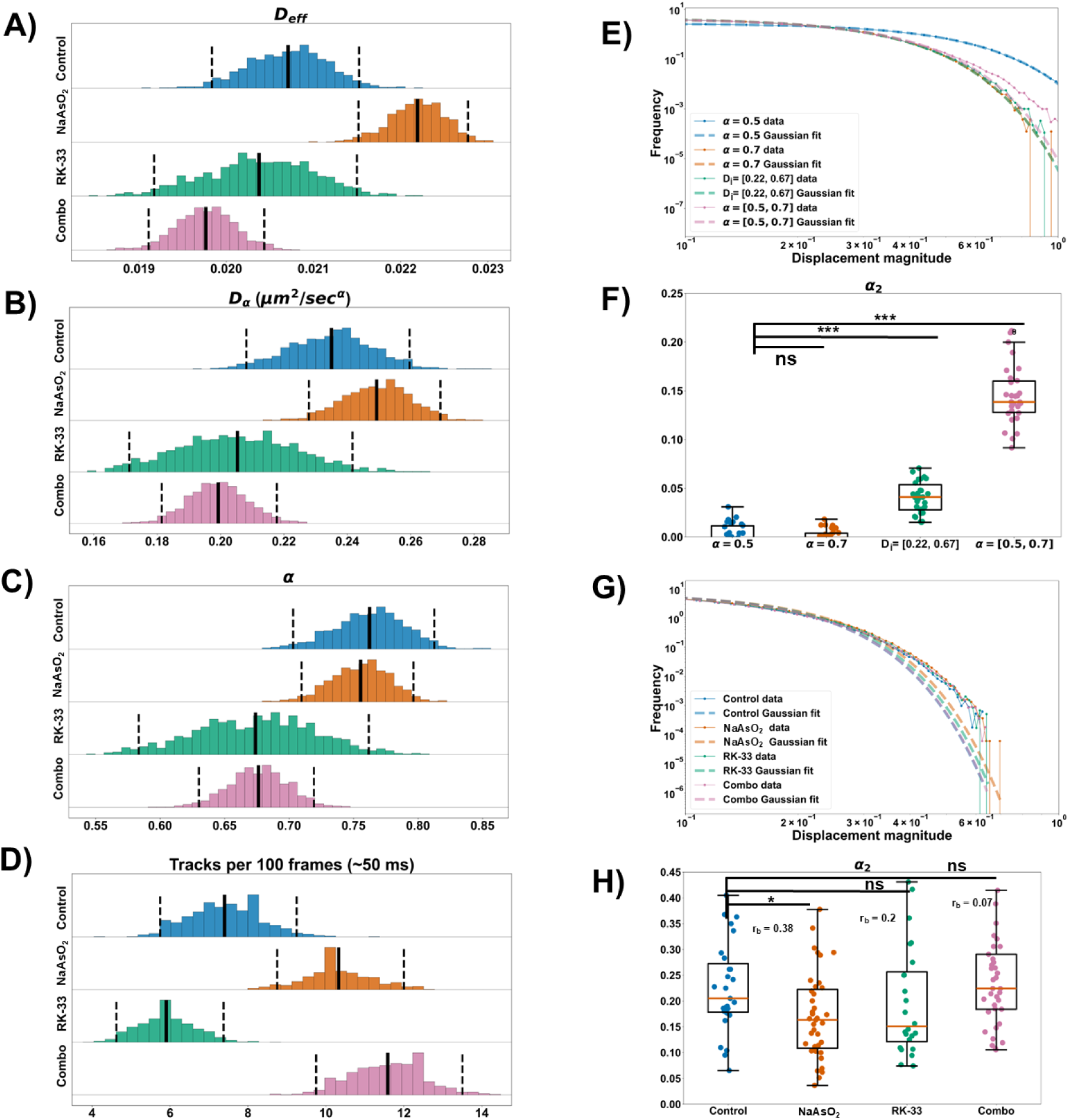
SPaCe-MC null distributions reveal cytoplasmic diffusion changes, while non-Gaussian displacement analysis quantifies cytoplasmic heterogeneity. **(A–D)** Null distributions of **(A)** Deff, **(B)** Dα, **(C)** α, and **(D)** track localization frequency generated by permutation of a circular mask (10 circles, area = 1.7 μm²) across segmented U2OS cells. Dashed black lines indicate the 2.5th and 97.5th percentiles (95% interval), and the solid line indicates the mean of each null distribution. Tracks within fixed circular masks, or within one permuted mask per cytoplasmic region, were pooled across videos. **(E, G)** Tails of track displacement distributions (10 ms lag time) shown on a log–log scale with Gaussian fits, and **(F, H)** the corresponding α₂. **(E, F)** Simulated cytoplasmic tracks with (i) α = 0.5, Di = 0.44 μm²/s^α^; (ii) α = 0.7, Di = 0.44 μm²/ s^α^; (iii) α = 0.7 and Di = 0.22–0.67 μm²/ s^α^; and (iv) Di = 0.44 μm²/ s^α^ and α = 0.5–0.7. **(G, H)** Experimental cytoplasmic tracks.

The resulting null distributions recapitulated the trends observed in the global cytoplasmic diffusion analysis (Fig. 1). NaAsO₂-treated cells showed increased D_eff_, with the entire 2.5–97.5 percentile range shifted toward higher values. While α remained largely unchanged, D_α_ shifted toward higher values, suggesting increased GEM mobility under similar anomalous scaling rather than a change in the mode of motion. This interpretation was further supported by simulations showing comparable D_α_ across tracks with identical α values (Figs. S17A–B and S18). In contrast, RK-33 alone or in combination with NaAsO₂ shifted both α and D_α_ toward lower values, indicating increased subdiffusive behavior.

Notably, the RK-33 null distributions were broader than those of the other treatments. To determine whether differences in null-distribution width reflected cytoplasmic diffusion heterogeneity or sampling statistics, we analyzed particle displacement distributions at a fixed 10-ms lag time and calculated the non-Gaussian parameter (α₂) (Fig. 5E–H; Fig. S2D–E; see Methods). Unlike the MSD-derived diffusion parameters described above, α₂ quantifies heterogeneity in particle displacement distributions rather than mean diffusion behavior. Consistent with previous reports^26^, simulations confirmed that α₂ is sensitive to heterogeneity in both α and D_α_ distributions (Fig. 5E–F; Fig. S17C–D). Experimentally, NaAsO₂ and RK-33 tended to decrease dynamic heterogeneity, reaching statistical significance for NaAsO₂, whereas the combined treatment did not differ significantly from control (Fig. 5G–H).

Altogether, these findings show that SPaCe-MC null distributions capture treatment-dependent changes in global cytoplasmic diffusion. Although dynamic heterogeneity also varied across SG-inducing conditions, differences in null-distribution width were not primarily driven by this heterogeneity. Instead, simulations indicated that the broader distributions observed in RK-33-treated cells were largely attributable to the lower number of recovered tracks (Figs. S3B and S18), demonstrating that null-distribution width is primarily determined by sampling density.

### Stress granule abundance and composition vary across SG-inducing conditions

To determine whether the SG-inducing conditions used in this study generated distinct SG populations, we quantified SG abundance, size, and the SG-associated proteins G3BP1 and DDX3. DDX3 was of particular interest because previous work showed that RK-33 alters its recruitment to SGs and changes RNA partitioning during SG formation^18,27^.

We confirmed that RK-33 treatment, either alone or prior to NaAsO₂ exposure, reduced SG abundance compared with NaAsO₂ treatment alone (Fig. 6C). RK-33-containing conditions also showed lower mean DDX3 fluorescence intensity within SGs (Fig. 6E), whereas SG volume decreased only with RK-33 alone. Comparison of U2OS cell lines with and without G3BP1-mCherry further showed that fluorescent G3BP1 expression influenced SG properties (Figs. S19–S21). Nevertheless, treatment-dependent differences in SG abundance and DDX3 recruitment were preserved in both cell lines, although the DDX3/G3BP1 intensity ratio depended on G3BP1-mCherry expression. Altogether, these results demonstrate that the SG-inducing conditions used in this study generate populations that differ not only in SG composition but also in SG abundance.

**Figure 6.**
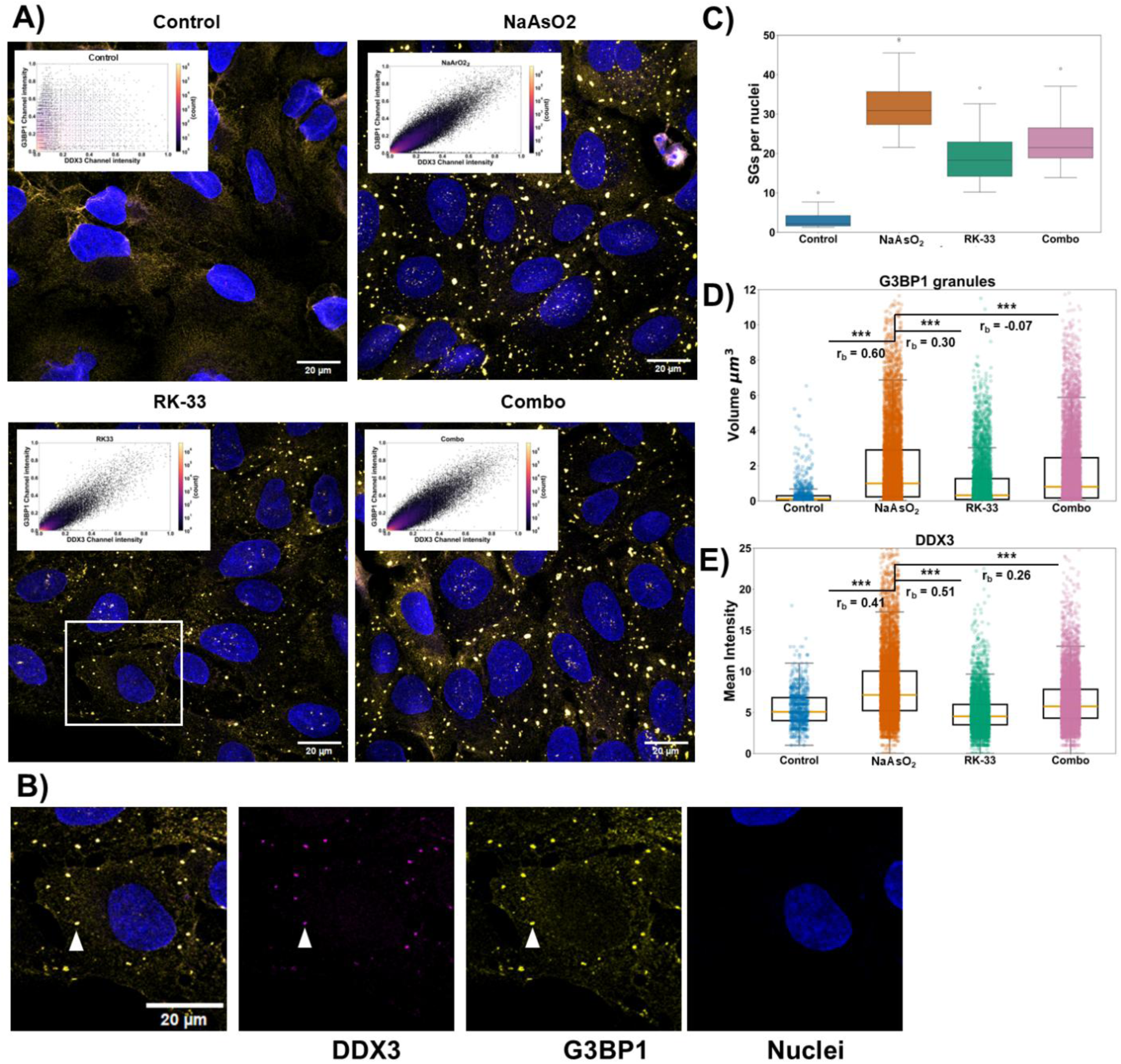
Changes in SG abundance and DDX3 enrichment in SGs under RK-33 treatment. **(A)** Maximum-intensity z-projections of U2OS cells and corresponding normalized cytofluorograms. DDX3 (magenta), G3BP1 (yellow), and nuclei (blue). Merged images are shown. **(B)** Cropped single-cell images showing merged and individual fluorescence channels. White arrows indicate G3BP1/DDX3-positive stress granules. **(C–E)** Quantitative analysis of SGs based on 3D segmentation of G3BP1-labeled z-stacks. **(C)** Number of SGs per nucleus. **(D)** SG volume. **(E)** Mean DDX3 fluorescence intensity within G3BP1-defined SGs. Each dot represents one SG.

## Discussion

In this study, we developed SPaCe-MC, an ROI-based statistical framework for distinguishing region-specific changes in intracellular diffusion from stochastic fluctuations and spatial sampling variability (Fig. 2). By comparing diffusion within predefined regions with treatment-matched, randomly sampled cytoplasmic regions, SPaCe-MC enables population-level testing of local diffusion changes in heterogeneous cellular environments.

Simulation-based validation identified sampling density and matching between observed and null distributions as key determinants of SPaCe-MC performance. Statistical power increased with track number and effect size, whereas mismatched sampling increased the FPR, specifically when the observed distribution was broader than the null (Fig. 3; Figs. S4–S15). Under sufficiently sampled, matched-null conditions, SPaCe-MC maintained an FPR close to the expected level while retaining high power to detect moderate changes in diffusion parameters. These findings establish practical constraints for applying SPaCe-MC and support population-level analysis when individual cellular regions contain insufficient tracks for reliable inference.

Having established these statistical properties, we applied SPaCe-MC to investigate cytoplasmic diffusion at the 40-nm length scale during SG formation. Our results reveal two distinct levels of cytoplasmic reorganization: global cytoplasmic changes and changes local to SGs.

Bulk cytoplasmic dynamics depended strongly on the SG-inducing condition (Figs. 1 and 5). NaAsO₂ increased GEM mobility, consistent with previous reports of stress-induced cytoplasmic fluidization^20^, with an increase in D_α_ but little change in the anomalous exponent α. In contrast, RK-33 treatment, either alone or in combination with NaAsO₂, increased the subdiffusive behavior of GEM nanoparticles. Dynamic heterogeneity also varied among SG-inducing conditions, with a tendency toward reduced heterogeneity following NaAsO₂ and RK-33 treatment that reached statistical significance for NaAsO₂ (Fig. 5G–H). Our previous work suggests that RK-33 alters RNA partitioning between the cytoplasm and ribonucleoprotein granule phases^27^, potentially contributing to changes in cytoplasmic diffusion through alterations in the RNA landscape via inhibition of DDX3’s normal role in RNA remodeling and helicase-dependent granule dynamics^23,24^. However, RK-33’s broader role in the stress response remains unclear, and although it is widely used as a DDX3 inhibitor^23,28–31^, RK-33 treatment itself is associated with induction of a stress response, potentially through DDX3-independent pathways^25^. While further experiments are required to dissect this effect, SG-inducing conditions appear to produce distinct global cytoplasmic diffusion responses that may obscure local, SG-associated diffusion changes.

Despite these differences in both bulk cytoplasmic dynamics and SG organization, including changes in SG abundance and DDX3 enrichment under RK-33-containing conditions, SPaCe-MC detected increased subdiffusive behavior within SG-associated regions across all treatments, as reflected by decreases in α and Dα (Fig. 4B–C). Particle localization frequency within SG-associated regions was generally comparable to that expected from the surrounding cytoplasm, except under NaAsO₂ treatment, where GEMs were more strongly excluded from SG regions (Fig. 4D; Fig. S3B–C). Super-resolution imaging showed that SG-associated GEMs localized predominantly near G3BP1-defined SG boundaries rather than within SG interiors, with most residing within ∼100 nm of the SG surface (Fig. 4F), consistent with the hypothesis of GEM exclusion from SGs^21^. The treatment-dependent difference in GEM exclusion may partly reflect differences in SG abundance, as the reduced SG abundance under RK-33-containing conditions (Fig. 6C) could limit the contribution of SG-associated exclusion to the overall cytoplasmic particle distribution. Thus, local constraints on GEM mobility can persist even when their contribution to the cytoplasmic-scale particle distribution is limited.

Collectively, integrating bulk cytoplasmic diffusion measurements with SPaCe-MC reveals spatially distinct physical responses within the cytoplasm. While the bulk cytoplasm exhibits inducer-dependent changes in nanoparticle mobility, SG-associated subcellular regions show a consistent local reduction in GEM mobility relative to their treatment-specific cytoplasmic background. These regions are spatially limited: cells contained approximately 20–30 SGs per nucleus with a median volume of ∼1 μm³ (Fig. 6C–D), corresponding to only ∼0.5–0.75% of total cellular volume based on an average U2OS cell volume^32^ of ∼4,000 μm³. Using a separate 2D segmentation-based measure, G3BP1-defined SG regions occupied ∼2–4% of the cytoplasmic area (Fig. S3A). Both estimates indicate that SG-associated regions represent a small fraction of the cellular space. Thus, spatially restricted SG-associated regions may contribute little to global diffusion measurements, while SPaCe-MC enables their behavior to be resolved relative to the surrounding cytoplasmic background.

SPaCe-MC may therefore provide a general framework for testing whether intracellular structures define local subcellular domains with diffusion properties distinct from those of the surrounding environment. Extending this approach to other organelles (for example, chromatin for nuclear diffusion^33^, etc.), cytoplasmic condensates and other structures (P-bodies, Q-bodies^20^, etc.), or membranes (lipid domains^34^, etc.) could reveal local physical heterogeneity that is obscured by cell-wide measurements. Such spatially resolved effects may be particularly relevant in polarized cells (such as neurons^35^) or multinuclear cells (fused macrophages^36^, placental cells^37^, etc.), where distinct subcellular regions may support specialized cellular functions.

## Acknowledgements

J.L.T. is the incipient SC SmartState Chair in Childhood Neurotherapeutics at the University of South Carolina. The authors gratefully acknowledge the computational resources provided by the Hyperion high-performance computing cluster at the University of South Carolina. We also acknowledge the technical assistance and resources provided by Research Computing at the University of South Carolina.

## Author Contributions

E.K., M.S., and P.V. designed the research. E.K., V.S., and P.V. performed the research and data analysis. E.K., P.V., and M.S. analyzed and interpreted the results. E.K., M.S., J.T., V.S., and P.V. wrote the manuscript. All authors have read and approved the manuscript.

## Data Availability

Microscopy data supporting this study are available on Figshare at https://doi.org/10.6084/m9.figshare.33860761 and https://doi.org/10.6084/m9.figshare.33858445. The deposited datasets include representative raw live-cell fluorescence time-lapse image series, stress-granule images, particle trajectories, and stress-granule segmentation masks.

## Methods

### Cell Culture and Treatment

U2OS cells stably expressing G3BP1-mCherry were kindly provided by the laboratory of Jeff Twiss. Tet-On doxycycline-inducible GEM-expressing cell lines were generated by transducing U2OS cells with the GEM construct (Addgene #231957) as described previously^11^. GEM expression was induced with 1 µg/mL doxycycline for 48 h.

For SG induction, cells were treated with 6 µM RK-33 in 0.15% DMSO or 0.15% DMSO alone for 2 h, with 500 µM NaAsO₂ added during the second hour.

Cells were maintained in high-glucose DMEM (ATCC) supplemented with 10% fetal bovine serum (Mediatech) and 1% penicillin–streptomycin (HyClone) at 37 °C and 5% CO₂.

### Live-Cell Imaging for Single-Particle Tracking

U2OS cells expressing doxycycline-inducible GEM (Addgene #231957) and G3BP1-mCherry were plated on black 0.17 mm Delta T dishes (04200417B). GEM expression was induced with 1 µg/mL doxycycline for 48 h before imaging. Live-cell imaging was performed on a Zeiss LSM 700 microscope equipped with a Delta T5 heated stage (35 °C) and a 100× oil-immersion objective. GEM fluorescence was acquired at 493/517 nm using an Axiocam 702 mono camera at maximum acquisition speed (200 fps, 5 ms exposure, ∼6 ms frame interval). G3BP1-mCherry images were acquired before GEM imaging at 540/590 nm (200 ms exposure). Images were collected at 512 × 512 pixels (75 × 75 µm field of view).

For each treatment condition, experiments were performed using two independently imaged cell populations (biological replicates), with approximately 15–20 videos acquired per replicate.

### Single-Particle Tracking, Track Pre-processing, and SG Segmentation

SPT was performed in Imaris using the Spots module. Particles were detected with an estimated diameter of 1 µm and background subtraction enabled and tracked using the Brownian motion algorithm with a maximum linking distance of 1 µm and a maximum gap of one frame. Spots with a quality score <3 and tracks shorter than 10 frames were excluded.

SG segmentation was performed using a custom Python pipeline implemented with SciPy and scikit-image. Images were normalized, and a cytoplasmic mask was generated by Gaussian smoothing (σ = 20) followed by intensity thresholding. Candidate SGs were identified within the cytoplasm using adaptive local thresholding (block size = 401 pixels) and Laplacian-of-Gaussian filtering (σ = 4), retaining pixels above the 95th percentile of the LoG response. Binary masks were hole-filled, labeled, and filtered by local contrast using the median intensity ratio between each SG and an expanded surrounding ring. Low-contrast and near-background objects were discarded. Final masks were saved as two-channel TIFF files containing SG and cytoplasmic masks.

### Extraction of Diffusion Parameters

To estimate bulk cytoplasmic diffusion parameters, all cytoplasmic particle tracks within each video were pooled for analysis. SG-associated local diffusion parameters were estimated from pooled tracks or track segments colocalizing with fixed SG masks across cells within each SG-inducing treatment. For each permutation, diffusion parameters were estimated from pooled tracks or track segments colocalizing with shuffled SG masks across cells within the corresponding SG-inducing treatment. Only tracks ≥10 frames (∼50 ms) were included.

Diffusion analysis followed our previous pipeline with minor modifications^11^. Briefly, the time- and ensemble-averaged mean squared displacement (MSD) was calculated from the x- and y-coordinates of all selected tracks. The first 100 ms of the MSD–lag time (τ) curve were fitted to the anomalous Brownian motion model:

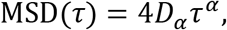

where *D_α_* is the anomalous diffusion coefficient and *α* is the anomalous diffusion exponent. D_eff_ was estimated from D_α_ and α as described previously^11^. The apparent Brownian diffusion coefficient (*D*) was estimated by fitting the first 50 ms of the MSD curve to the Brownian diffusion model (*α* = 1). A minimum lag time of 10 ms was used for all fits.

### Spatially Constrained Monte Carlo Permutation Test (SPaCe-MC)

For SPaCe-MC analysis, the observed statistic was calculated from particle tracks within SG masks pooled across all videos for a given treatment. The null distribution was generated by randomly relocating SG masks throughout the cytoplasm while excluding existing SG regions. Tracks contained within the permuted masks were pooled across cells to estimate the corresponding diffusion parameter. This procedure was repeated 1,000 times to generate the null distribution.

The 95% confidence interval was defined by the 2.5th and 97.5th percentiles of the null distribution. Statistical significance was assessed using a two-sided permutation test by comparing the absolute deviation of the observed statistic from the null mean with the distribution of absolute deviations of the permuted statistics. The *p*-value was calculated as the proportion of permuted statistics with deviations at least as large as the observed value.

### Analysis of Non-Gaussian Displacement Statistics

To assess dynamic heterogeneity beyond mean diffusion parameters, we quantified the non-Gaussian parameter (α₂) from particle displacement distributions. For each track, x-and y-displacements were calculated at a lag time of 10 ms and pooled within each treatment group to generate a self-displacement distribution, often referred to as the self-part of the Van Hove distribution. The non-Gaussian parameter (α₂) was calculated as described previously^26^:

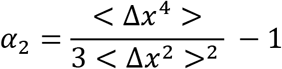

To visualize deviations from Gaussian diffusion, Van Hove distributions were plotted on a log– log scale.

### Simulation of Fractional Brownian Motion

Particle tracks were simulated using the Python FBM package, which generates fractional Brownian motion (fBM) tracks with a specified Hurst exponent (H). Independent x- and y-tracks were generated and scaled to the desired input diffusion coefficient (D_i_).

To reproduce the sampling characteristics of each treatment, we generated 1,000 simulated track files, each containing track groups matching the number of experimentally analyzed videos (Fig. S3). Null datasets used the median track number and duration across experimental permutations (Fig. S3C). Two observation sampling schemes were evaluated: (i) matched sampling, in which track number and duration were drawn from triangular distributions matching the null statistics (Figs. S4–S9), and (ii) experimental sampling, in which observation datasets matched the SG-associated track statistics (Fig. S3B) while null datasets retained the permutation-derived sampling characteristics (Figs. S10–S15).

Null tracks were simulated with H = 0.35 (α = 0.70) and D_i_ = 0.45 μm²/s. Observation datasets were generated by varying either H (0.25–0.35) at fixed D_i_ = 0.45 μm²/s or D_i_ (0.27–0.44 μm²/s) at fixed H = 0.35.

Diffusion parameters were estimated as described above, and a two-sided permutation-test *p*-value was calculated for each simulation. Resulting *p*-value distributions were used to estimate the false-positive rate (FPR), false-negative rate (FNR), and statistical power. FPR was estimated using simulations with identical observed and null parameters (α = 0.70), whereas power was estimated using progressively lower observed α values (0.65–0.40). Statistical significance was defined as *p* < 0.05.

To evaluate the effects of sampling and diffusion parameters on SPaCe-MC null distributions, we generated simulated datasets consisting of 30 track groups (30 videos) while systematically varying H, D_i_, track number, or track duration (Fig. S16). Diffusion parameters and α₂ were estimated as described above (Fig. S17). To examine the effect of sampling density, diffusion parameters were calculated from random subsets of 20, 250, or 1,000 tracks per permutation, representing low-(single-video), intermediate-(RK-33), and high-sampling (NaAsO₂ and Combo) conditions (Fig. S18).

### Immunocytochemistry

U2OS cells were grown on 18 mm Deckglaser GG-18-1.5H coverslips in 12-well plates. At 60– 80% confluency, cells were treated with NaAsO₂ with or without RK-33 as described above, fixed with 4% paraformaldehyde for 10 min, permeabilized with 0.1% Triton X-100 in PBS for 15 min, and blocked in 10% FBS or BSA in PBS for 1 h at room temperature. Primary antibodies diluted in blocking buffer were incubated for 2 h at room temperature.

Primary antibodies were anti-G3BP1 (rabbit, 1:750; A302-033A, Thermo Fisher Scientific) and anti-DDX3 (mouse, 1:750; sc-365768, Santa Cruz Biotechnology). After three PBS washes, samples were incubated for 1.5 h with donkey anti-rabbit Alexa Fluor Plus 488 (10 µg/mL; A32790, Invitrogen) or STAR RED (5 µg/mL; STRED-1002-500UG, Abberior), and anti-mouse CF640R (10 µg/mL; 20175, Biotium) or STAR ORANGE (5 µg/mL; STORANGE-1001-500UG, Abberior). Cells were counterstained with 1 µM DAPI for 10 min, washed with PBS, and mounted in ProLong Glass Antifade Mountant (P36980, Invitrogen) on Superfrost slides.

### Collection of Confocal and STED Z-stacks

Confocal and STED imaging were performed on a Stellaris 8 STED 3D FALCON microscope (Leica Microsystems) equipped with an HC PL APO CS2 100×/1.40 NA oil-immersion objective (pinhole 151.6 µm).

Confocal z-stacks were acquired using a scan speed of 200 Hz, 0.75× zoom, 0.30 × 0.30 × 0.23 µm voxels, and line averaging of 2. Alexa Fluor Plus 488 (G3BP1), CF640R (DDX3), and DAPI were excited at 587, 631, and 499 nm using laser powers of 4%, 4%, and 0.3%, respectively.

STED z-stacks were acquired at 400 Hz, 2× zoom, 0.11 × 0.11 × 0.10 µm voxels, and line averaging of 4. Sapphire was excited at 483 nm (4%), and STAR RED-labeled G3BP1 at 631 nm (5.52%) with depletion by a 775 nm STED laser (50.2%). Three-dimensional STED imaging used both the Vortex 780 (Beam 1) and z-phase 780 (Beam 2) phase masks.

For each treatment condition, experiments were performed using two independently imaged cell populations (biological replicates). Five confocal z-stacks (155 × 155 ηm XY field of view) and five STED z-stacks (58 × 58 ηm XY field of view) were acquired per replicate.

### Stress Granule Segmentation and Colocalization Analysis

Confocal SG z-stacks were segmented in Python using a Laplacian-of-Gaussian (LoG)-based pipeline. Z-slices were first filtered by the 90th-percentile intensity of channel 2, retaining slices with intensities ≥20% of the brightest slice. Nuclei were segmented from channel 3 using Gaussian smoothing, Otsu thresholding, hole filling, and small-object removal.

SGs were detected independently in each fluorescence channel using two-scale LoG filtering (σ = 2 and 1.5) followed by 96th-percentile intensity thresholding. Small objects overlapping larger granules were excluded, and binary masks were hole-filled, labeled in 3D, and filtered by a minimum maximum voxel intensity of 10.

Morphological measurements were extracted using regionprops and included volume, centroid, mean and maximum intensity, bounding-box coordinates, and surface area estimated by marching-cubes reconstruction. SG and nuclear counts were also recorded. For colocalization analysis, per-object summed intensity and channel intensity ratios were calculated. Signal colocalization was visualized using min-max normalized voxel intensities, percentile-based background filtering, and hexagonal density plots.

For STED–confocal analysis, the same pipeline was applied, with GEM particles segmented using LoG filtering (σ = 2 and 1) and an intensity threshold of 1. Particle positions were represented by center-of-mass coordinates, and Euclidean distance transforms of calibrated SG masks were used to calculate signed distances to SG surfaces (negative inside, positive outside). SGs were subdivided into interior, peripheral, and neighboring regions using two erosions and three dilations; granules too small for subdivision were classified as small SGs. SG volumes were calculated from calibrated voxel counts.

### Statistical Analysis

Statistical analyses were performed using two-sided Mann–Whitney U tests for data presented as boxplots. Statistical significance was defined as ∗p < 0.05, ∗∗p < 0.01, ∗∗∗p < 0.001. Effect size was quantified using the rank-biserial correlation (rb), interpreted as small (rb < 0.3), medium (0.3 ≤ rb < 0.5), or large (rb ≥ 0.5)^11^.

## Code availability

The Python implementation of the SPaCe-MC framework is freely available at: https://github.com/Korunova/space_mc_paper

