## Supplemental Materials for "Spatially Constrained Monte Carlo Permutation Test Reveals Diffusion Changes Near Stress Granules"

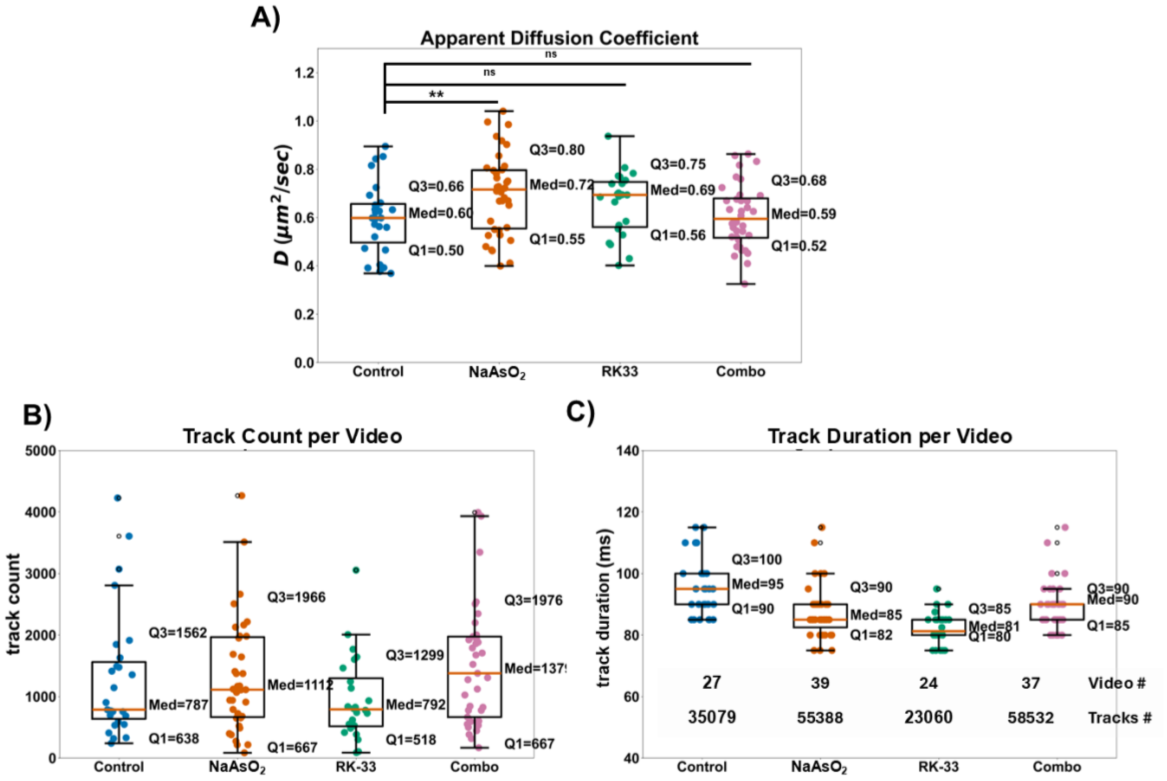


**Figure S1. Apparent Diffusion coefficient**

**(A)** Apparent diffusion coefficient (D) obtained from classical Brownian motion fits (R2 filter ≥ 0.7, 50 ms). Cells were imaged at the onset of $NaAsO_2$ treatment (1 h after RK-33 or DMSO treatment) and analyzed after 30 min of imaging, corresponding to robust $NaAsO_2$-induced SG formation. Each dot corresponds to the diffusion parameter estimated from all GEM trajectories detected within that one video (75 × 75 μm field of view).

Track parameters obtained from videos: **(B)** number of tracks per video and **(C)** track duration per video.

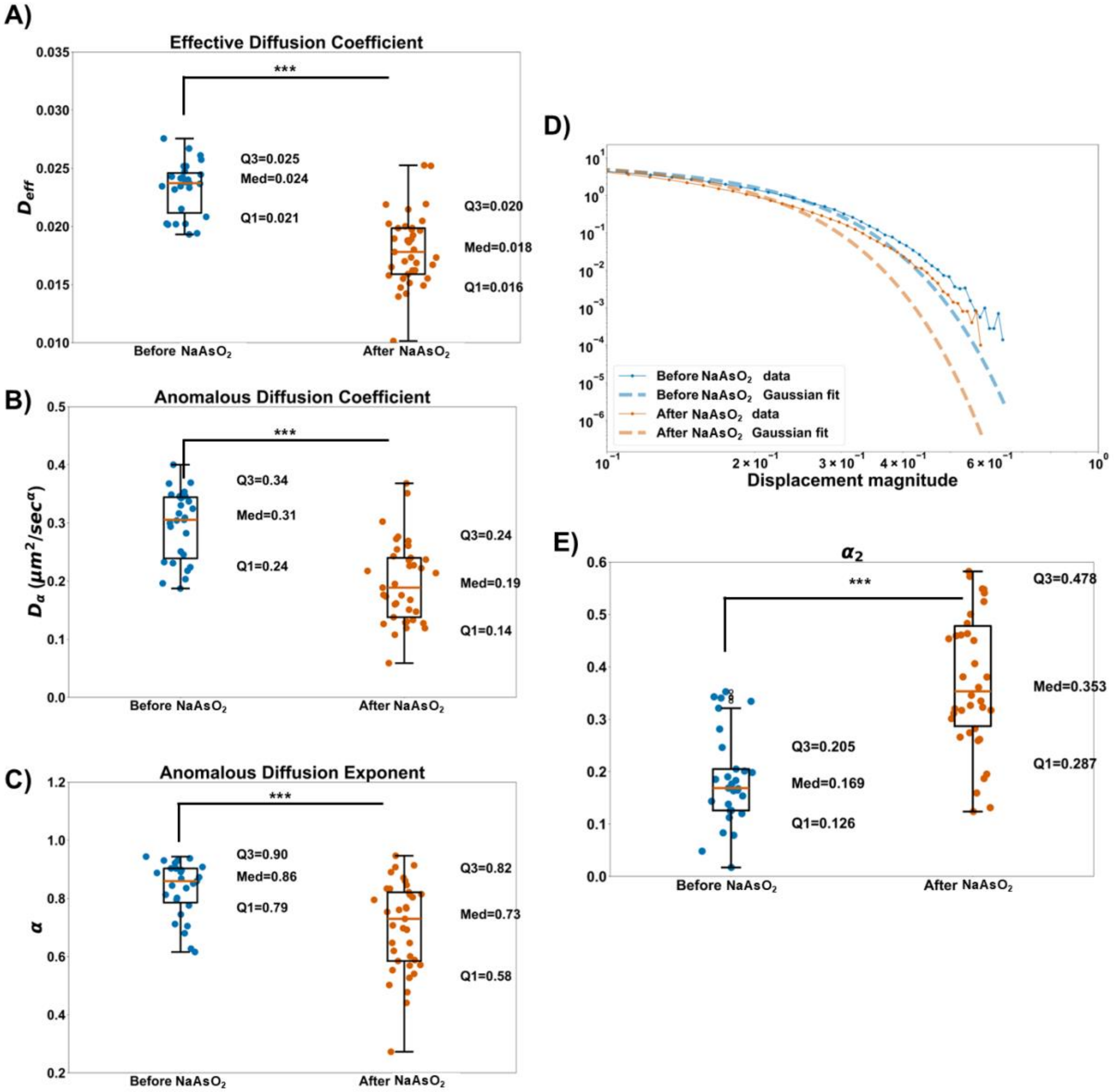


**Figure S2. Reduced Diffusivity and Increased Heterogeneity of Particle Motion During Repeated Imaging Before and After $NaAsO_2$ Treatment**

(A–C) Diffusion parameters extracted from anomalous Brownian motion fits ($R^2 > 0.9$, 100 ms): (A) effective diffusion coefficient ($D_{eff}$) (B) anomalous exponent (α) and (C) anomalous diffusion coefficient ($D_\alpha$).

(D) Tails of the GEM particle displacement distributions (10 ms lag time) shown on a log–log scale with Gaussian fits, and (E) the corresponding non-Gaussian parameter ($\alpha_2$) of the particle displacement distributions.

All estimated parameters were averaged over the cytoplasm in the video (75 x 75 μm field of view). Cells were imaged at the onset of $NaAsO_2$ treatment (1 h after RK-33 or DMSO treatment) and analyzed after immediately after imaging.

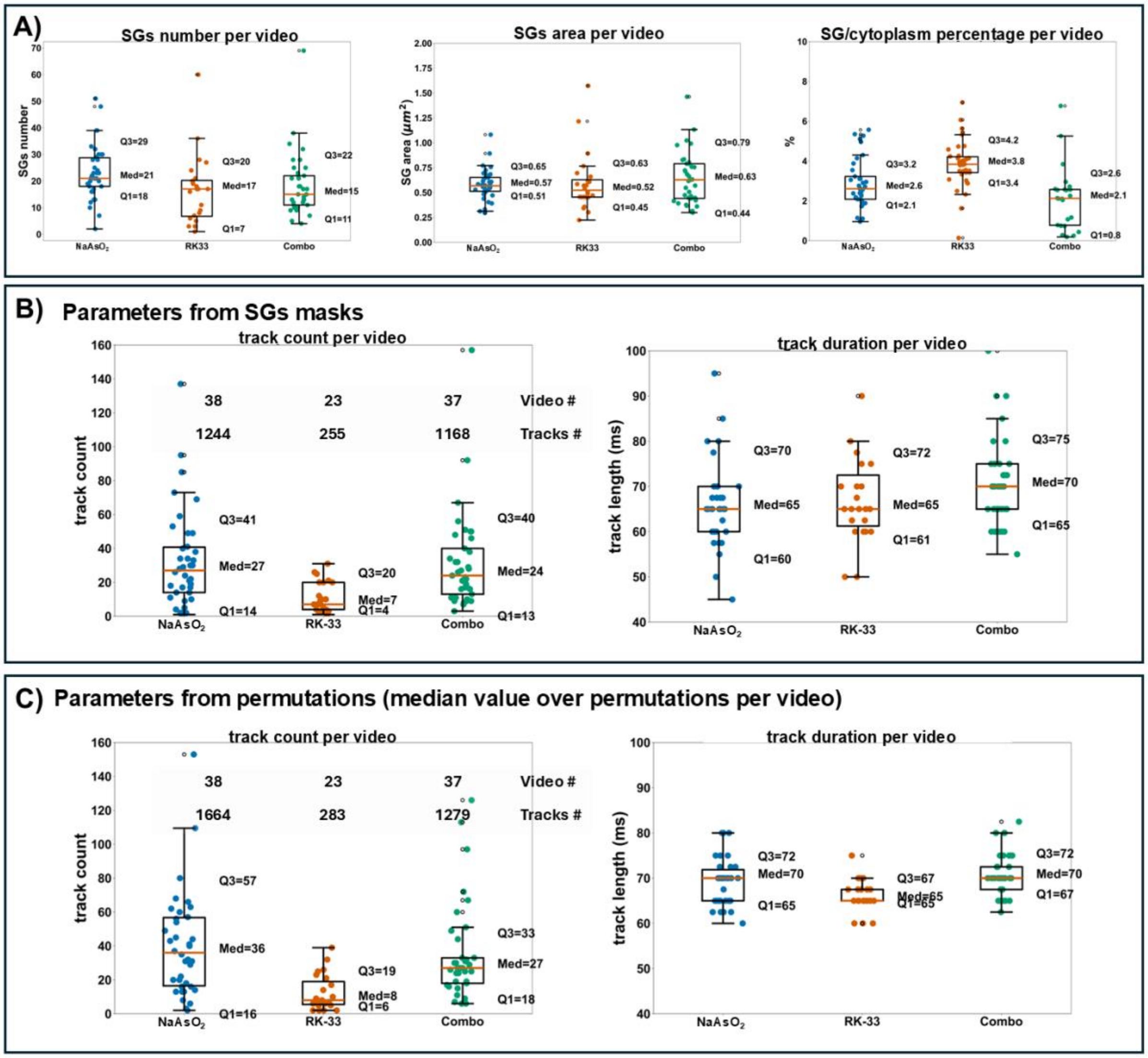


**Figure S3. Parameters of stress granules (SGs) and collected particle tracks.**

(A) SG parameters: (left) number of SGs per video, (middle) total SG area per video, and (right) percentage of SG area to cytoplasm area per video.

(B) Track parameters obtained from SG masks: (left) number of tracks per video and (right) track duration per video.

(C) Median track statistics across all permutations for each video: (left) number of tracks per video and (right) track duration per video.

One video – 400 frames (~6 ms per frame), 512 × 512 pixels (75 μm × 75 μm field of view).

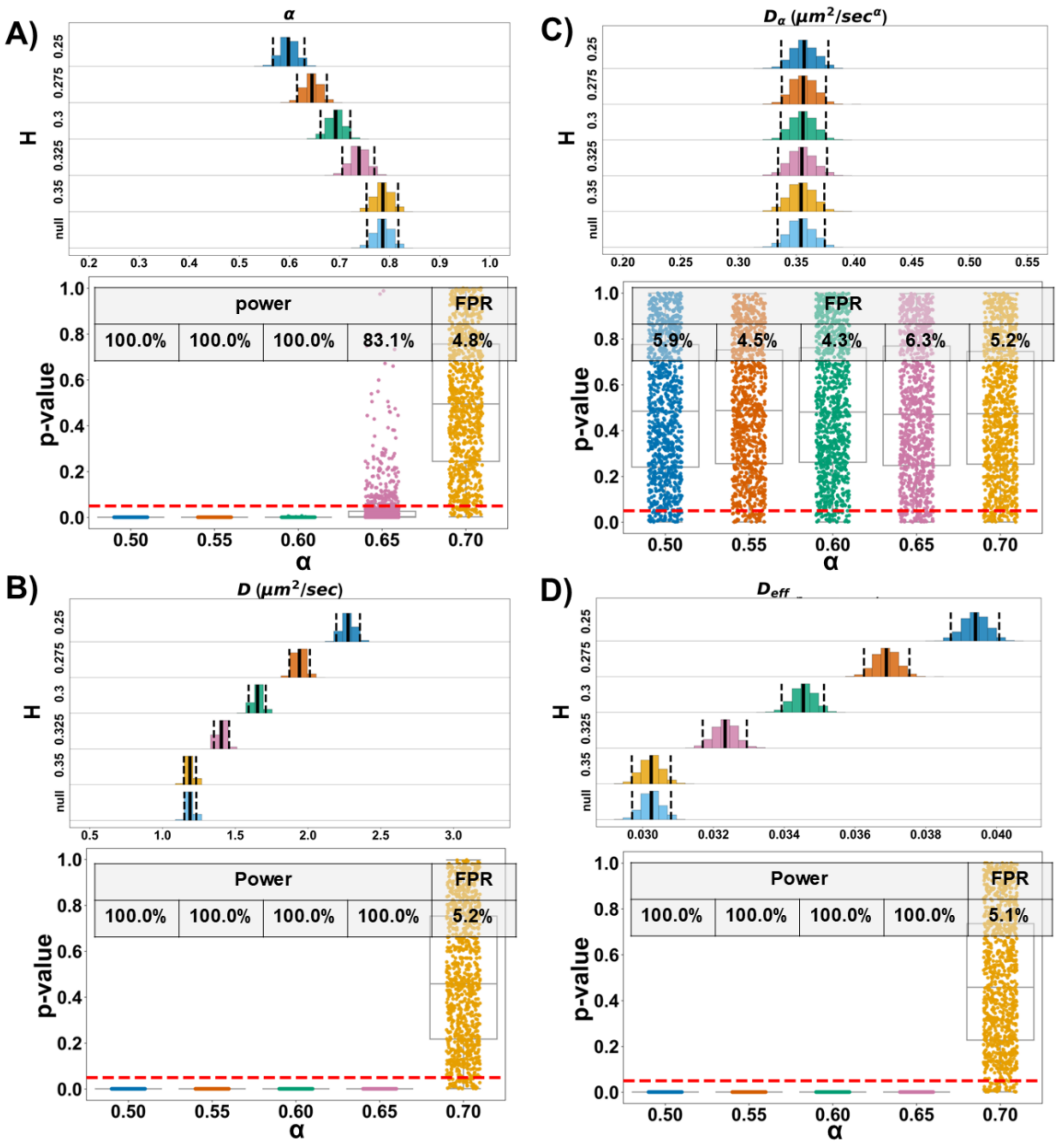


**Figure S4. Validation of the SPaCe-MC Permutation Framework Using Simulated Track Datasets with Observation Track Statistics Derived from Permutations During Combo Treatment: Variability in the Hurst Parameter Between Groups**

(A–D) Joyplots showing distributions of the diffusion parameters α (A), D (B), $D_{\alpha}$ (C) and $D_{eff}$ (D) together with boxplots of permutation-test p-values obtained from each simulation. Simulations were generated with Hurst parameters of 0.25, 0.30, 0.325, and 0.35 (corresponding to α = 0.50, 0.60, 0.65, and 0.70) while keeping the diffusion coefficient fixed at 0.44 $\mu m^2$/s.

Simulated datasets (both the null distribution and observations) matched the distributions of track number and track duration obtained from the permutation analysis (Combo; Fig. S3C) but did not preserve spatial organization. p-value was 0.05 for power and FPR estimates.

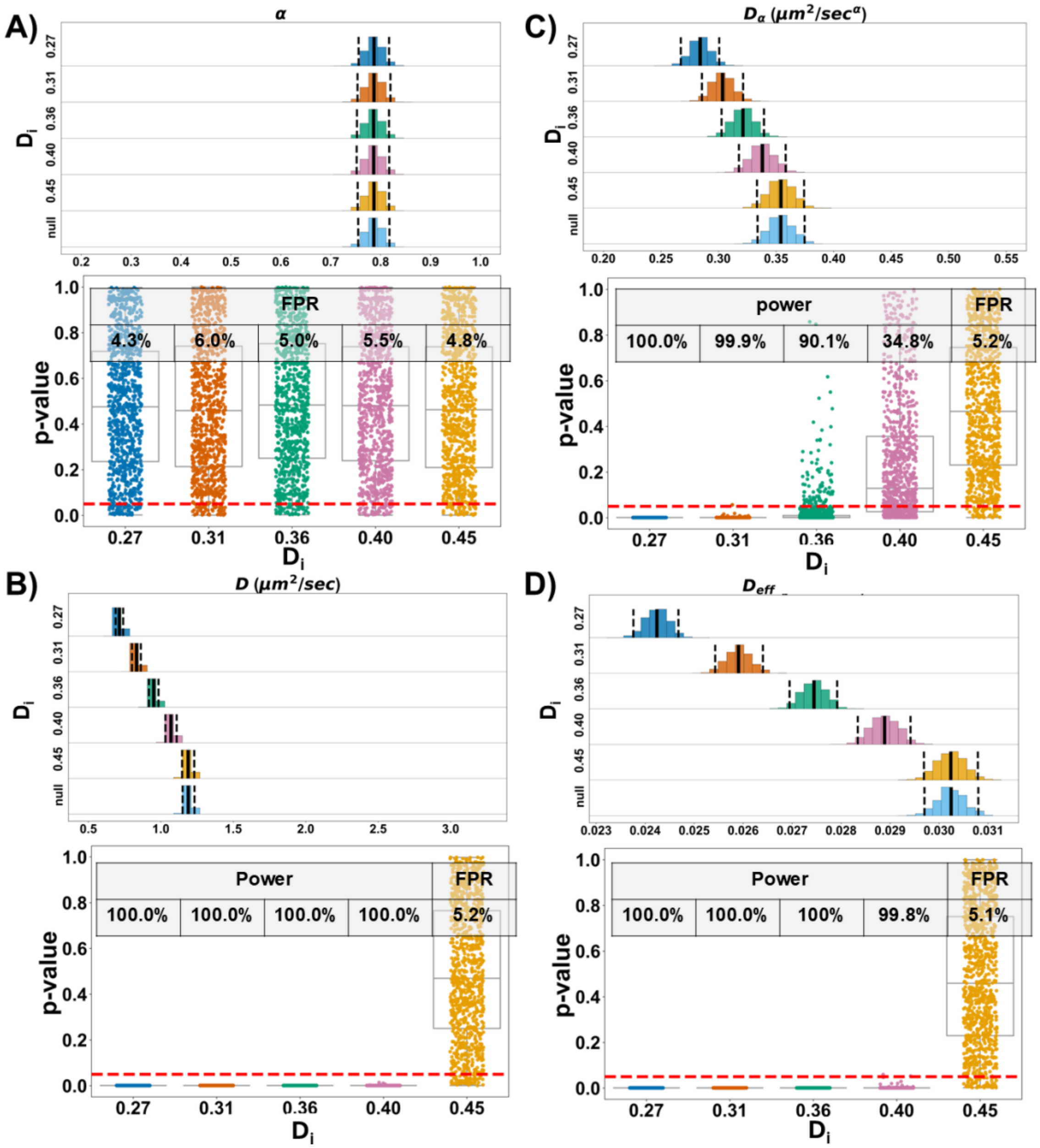


**Figure S5. Validation of the SPaCe-MC Permutation Framework Using Simulated Track Datasets with Observation Track Statistics Derived from Permutations During Combo Treatment: Variability in Diffusion Coefficient Between Groups**

(A–D) Joyplots showing distributions of the diffusion parameters α (A), D (B), $D_\alpha$ (C) and $D_{eff}$ (D) together with boxplots of permutation-test p-values obtained from each simulation. Simulations were generated with input diffusion coefficient ($D_i$) of 0.27, 0.31, 0.36, 0.40, and 0.44 $\mu m^2$/s while keeping the Hurst parameter fixed at 0.35.

Simulated datasets (both the null distribution and observations) matched the distributions of track number and track duration obtained from the permutation analysis (Combo; Fig.

S3C) but did not preserve spatial organization. p-value was 0.05 for power and FPR estimates.

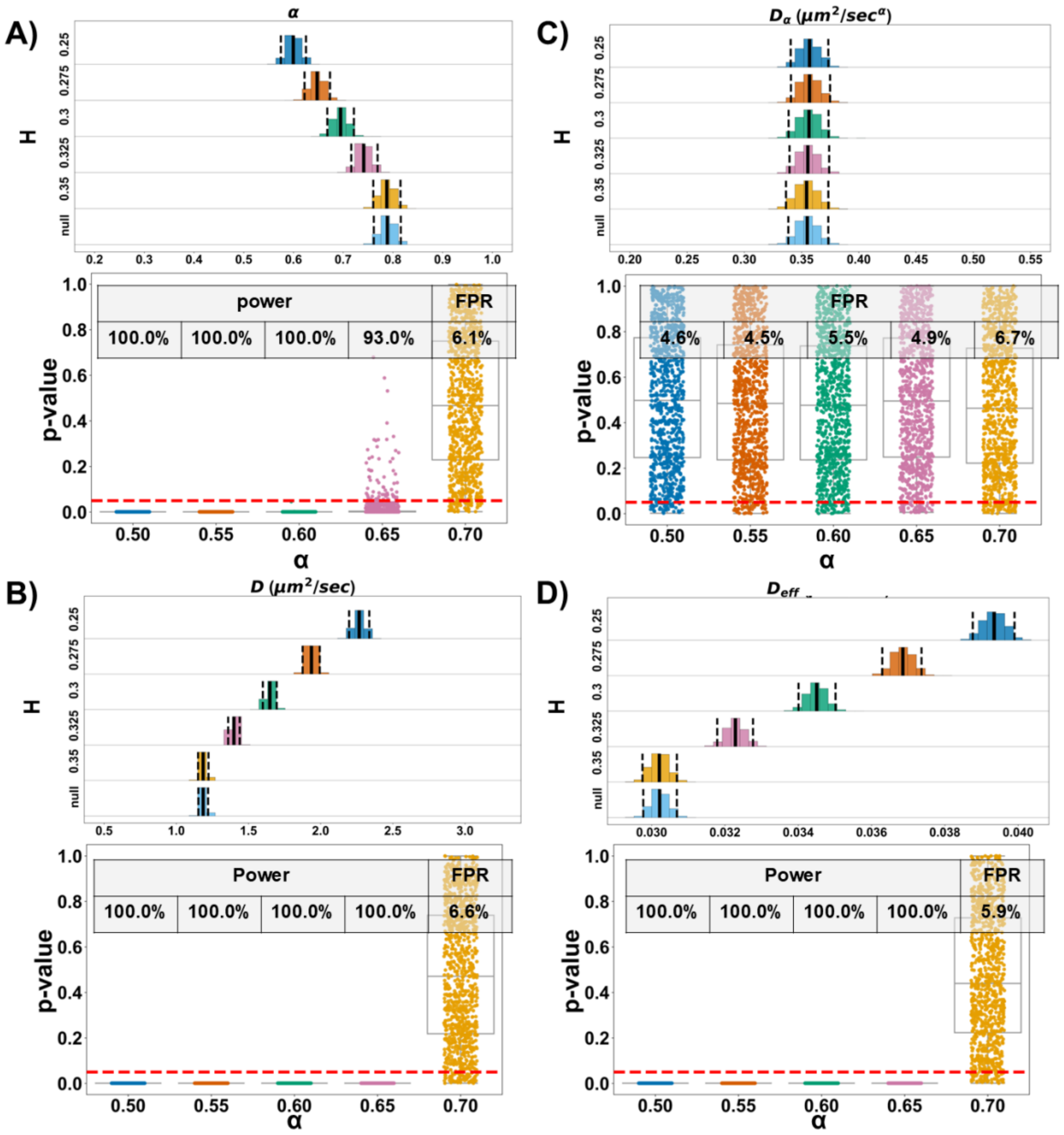


**Figure S6. Validation of the SPaCe-MC Permutation Framework Using Simulated Track Datasets with Observation Track Statistics Derived from Permutations During $NaAsO_2$ Treatment: Variability in the Hurst Parameter Between Groups**

(A–D) Joyplots showing distributions of the diffusion parameters α (A), D (B), $D_\alpha$ (C) and $D_{eff}$ (D) together with boxplots of permutation-test p-values obtained from each simulation. Simulations were generated with Hurst parameters of 0.25, 0.30, 0.325, and 0.35 (corresponding to α = 0.50, 0.60, 0.65, and 0.70) while keeping the diffusion coefficient fixed at 0.44 $\mu m^2/s$.

Simulated datasets (both the null distribution and observations) matched the distributions of track number and track duration obtained from the permutation analysis (Combo; Fig. S3C) but did not preserve spatial organization. p-value was 0.05 for power and FPR estimates.

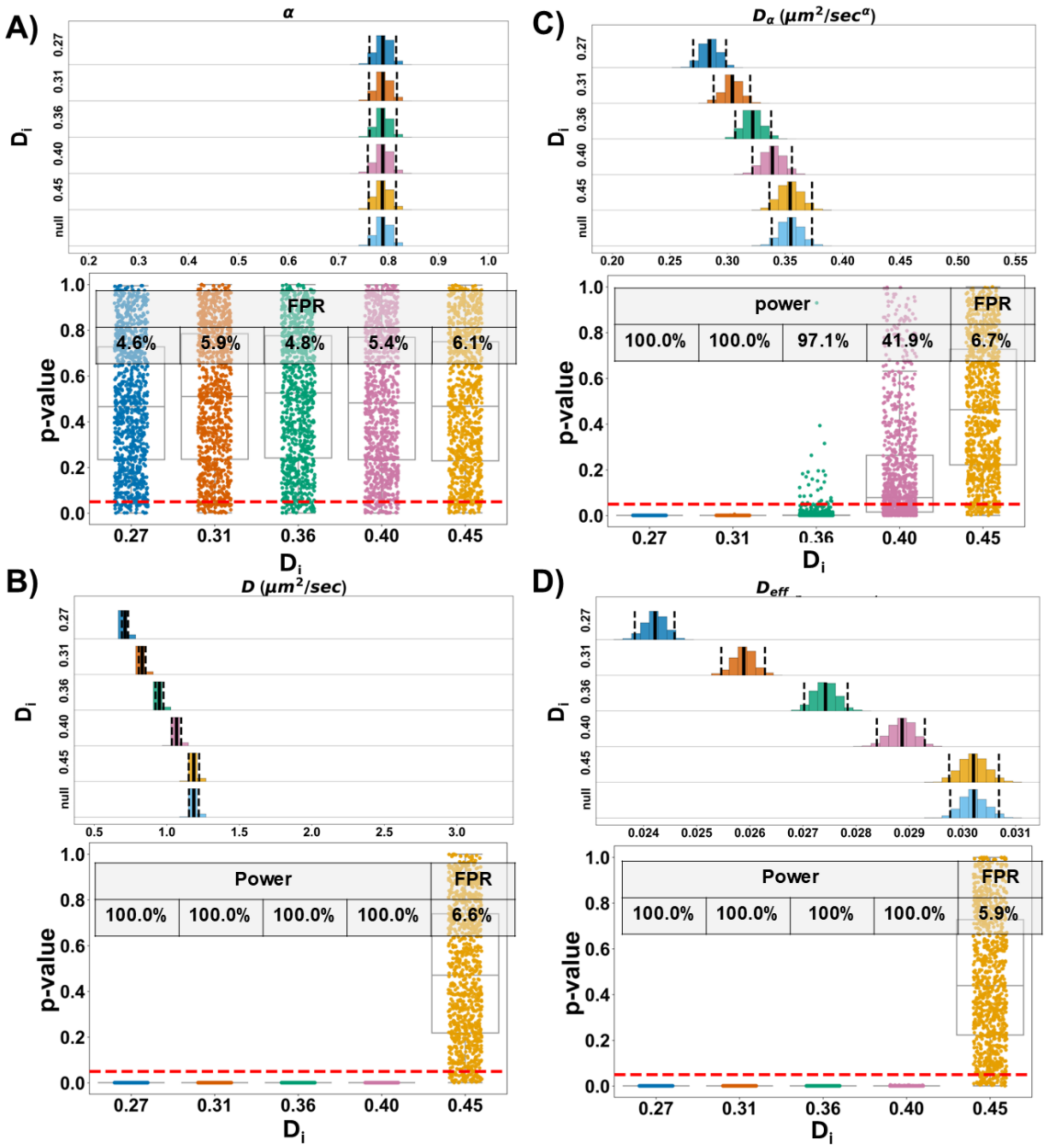


**Figure S7. Validation of the SPaCe-MC Permutation Framework Using Simulated Track Datasets with Observation Track Statistics Derived from Permutations During $NaAsO_2$ Treatment: Variability in Diffusion Coefficient Between Groups**

(A–D) Joyplots showing distributions of the diffusion parameters α (A), D (B), $D_\alpha$ (C) and $D_{eff}$ (D) together with boxplots of permutation-test p-values obtained from each simulation. Simulations were generated with input diffusion coefficient ($D_i$) of 0.27, 0.31, 0.36, 0.40, and 0.44 $\mu m^2/s$ while keeping the Hurst parameter fixed at 0.35.

Simulated datasets (both the null distribution and observations) matched the distributions of track number and track duration obtained from the permutation analysis (Combo; Fig.

S3C) but did not preserve spatial organization. p-value was 0.05 for power and FPR estimates.

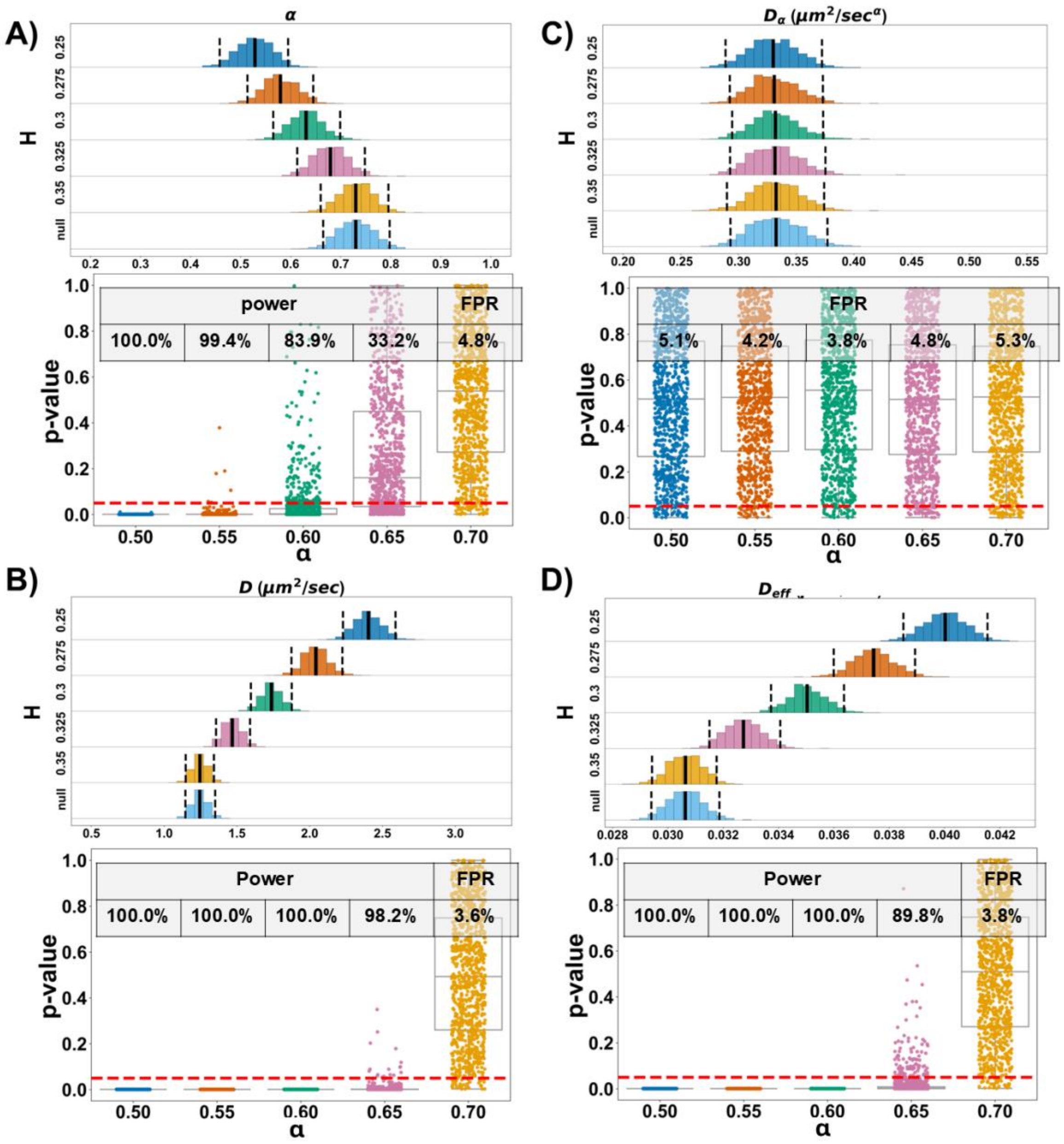


**Figure S8. Validation of the SPaCe-MC Permutation Framework Using Simulated Track Datasets with Observation Track Statistics Derived from Permutations During RK-33 Treatment: Variability in the Hurst Parameter Between Groups**

(A–D) Joyplots showing distributions of the diffusion parameters α (A), D (B), $D_\alpha$ (C) and $D_{eff}$ (D) together with boxplots of permutation-test p-values obtained from each simulation. Simulations were generated with Hurst parameters of 0.25, 0.30, 0.325, and 0.35 (corresponding to α = 0.50, 0.60, 0.65, and 0.70) while keeping the diffusion coefficient fixed at 0.44 $\mu m^2/s$.

Simulated datasets (both the null distribution and observations) matched the distributions of track number and track duration obtained from the permutation analysis (Combo; Fig. S3C) but did not preserve spatial organization. p-value was 0.05 for power and FPR estimates.

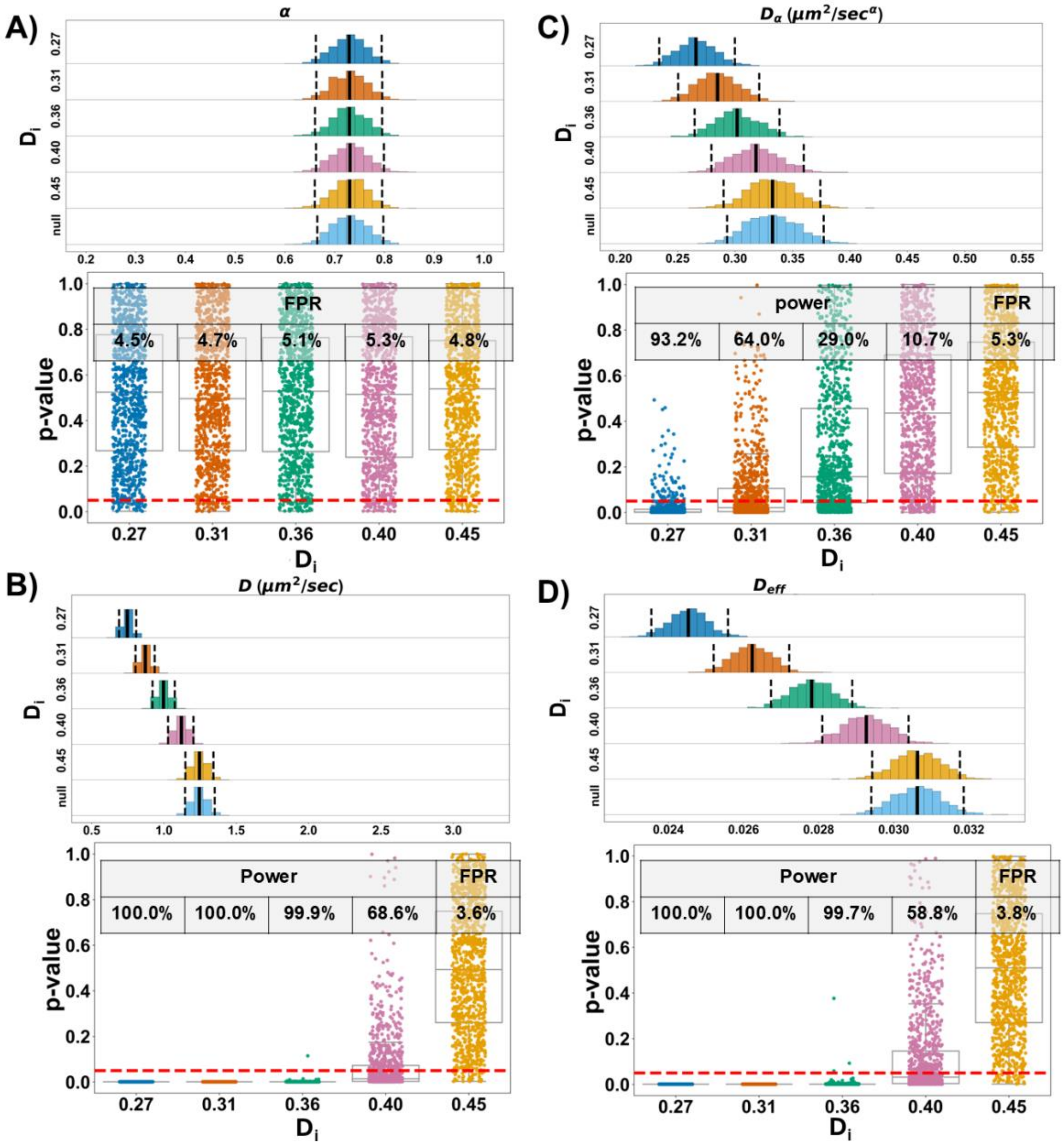


**Figure S9. Validation of the SPaCe-MC Permutation Framework Using Simulated Track Datasets with Observation Track Statistics Derived from Permutations During RK-33 Treatment: Variability in Diffusion Coefficient Between Groups**

(A–D) Joyplots showing distributions of the diffusion parameters α (A), D (B), $D_\alpha$ (C) and $D_{eff}$ (D) together with boxplots of permutation-test p-values obtained from each simulation. Simulations were generated with input diffusion coefficient ($D_i$) of 0.27, 0.31, 0.36, 0.40, and 0.44 μm$^2$/s while keeping the Hurst parameter fixed at 0.35.

Simulated datasets (both the null distribution and observations) matched the distributions of track number and track duration obtained from the permutation analysis (Combo; Fig.

S3C) but did not preserve spatial organization. p-value was 0.05 for power and FPR estimates.

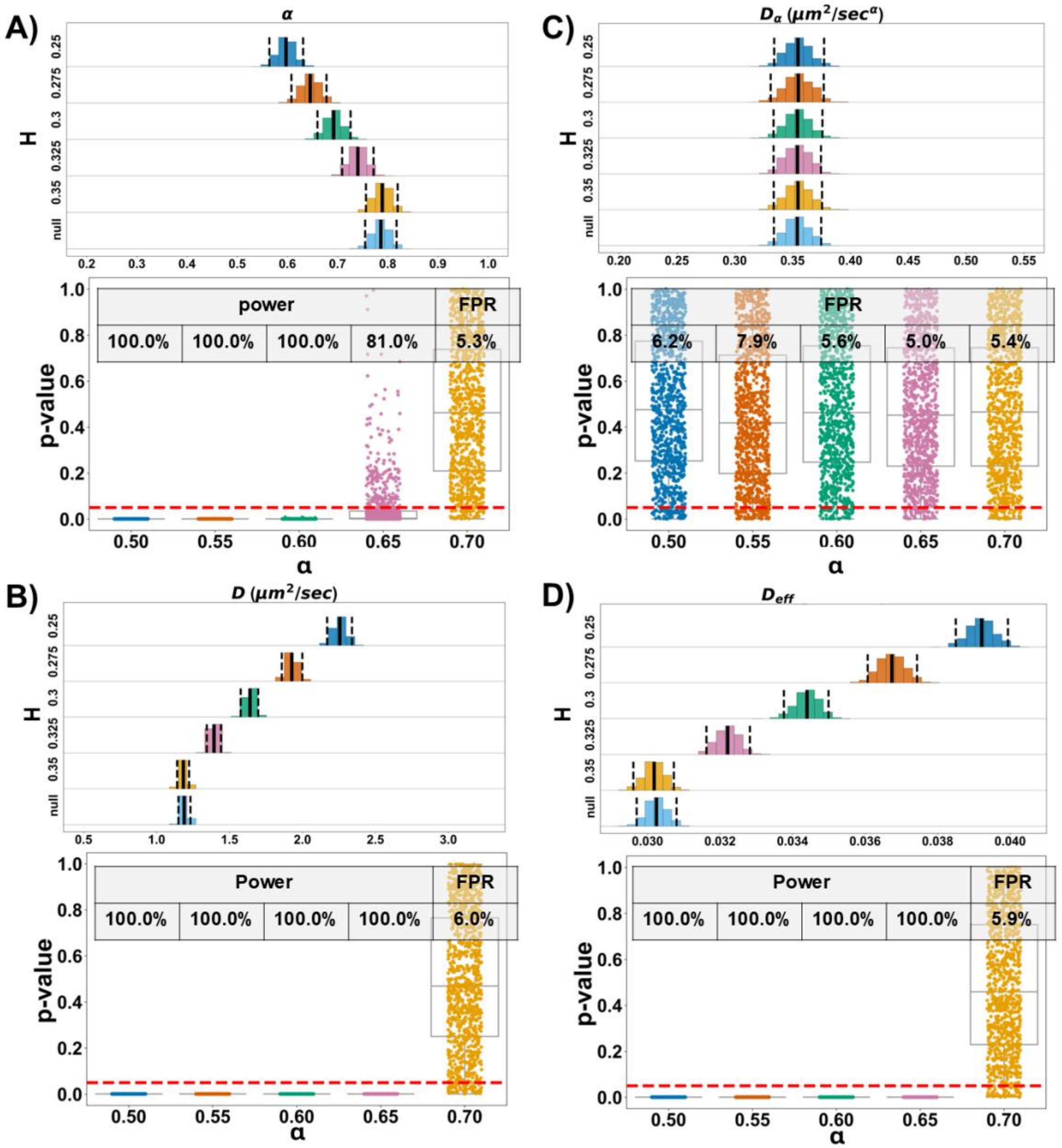


**Figure S10. Validation of the SPaCe-MC Permutation Framework Using Simulated Track Datasets with Experimental Track Statistics from Combo Treatment: Variability in the Hurst Parameter Between Groups**

(A–D) Joyplots showing distributions of the diffusion parameters α (A), D (B), $D_\alpha$ (C) and $D_{eff}$ (D) together with boxplots of permutation-test p-values obtained from each simulation. Simulations were generated with Hurst parameters of 0.25, 0.30, 0.325, and 0.35 (corresponding to α = 0.50, 0.60, 0.65, and 0.70) while keeping the diffusion coefficient fixed at 0.44 $\mu m^2/s$.

Null simulated datasets matched the distributions of track number and track duration obtained during the permutation analysis (Combo; Fig. S3C), whereas the simulated observation datasets matched the experimental sample size (Combo; Fig. S3B) but did not preserve spatial organization. p-value was 0.05 for power and FPR estimates.

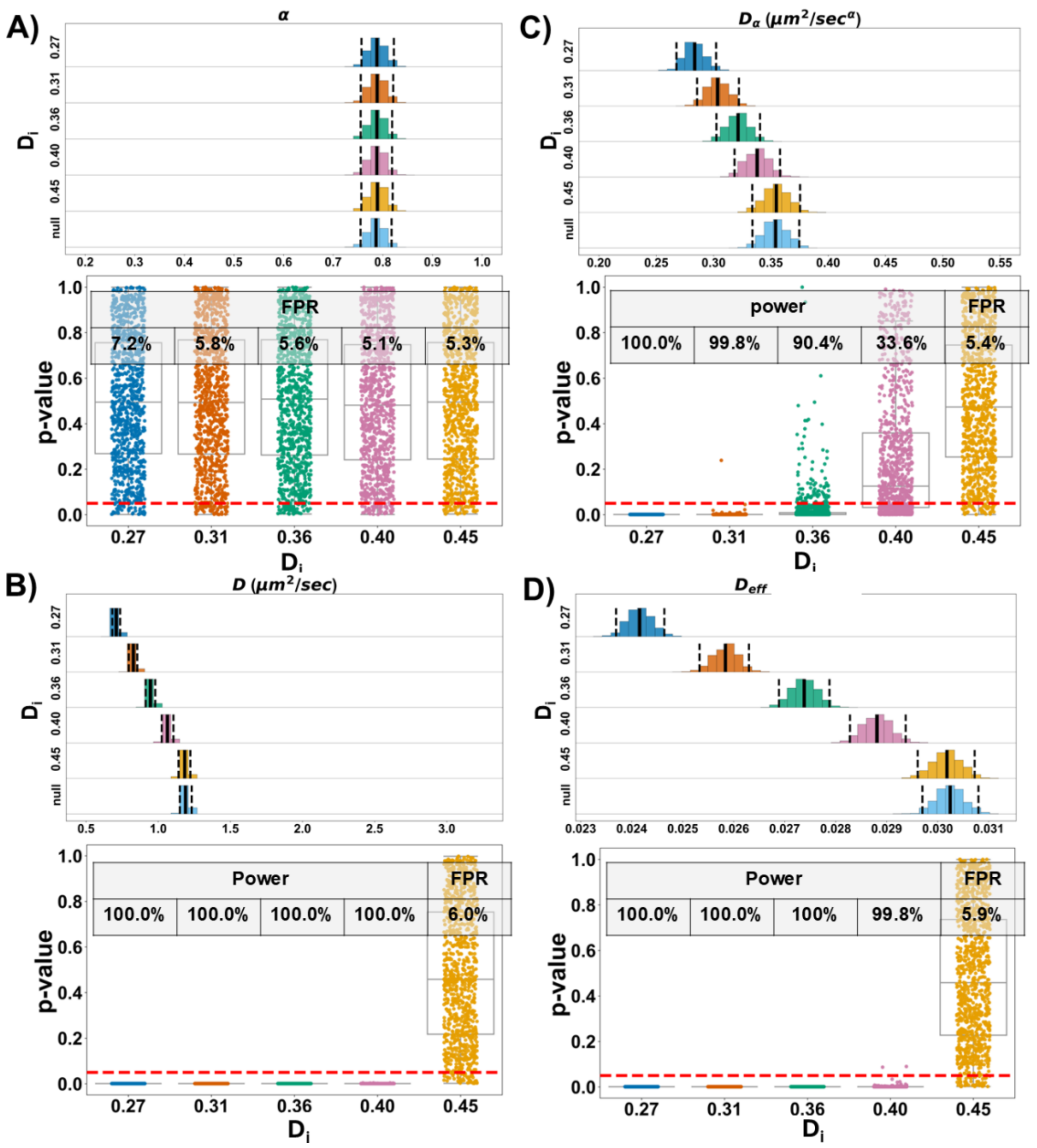


**Figure S11. Validation of the SPaCe-MC Permutation Framework Using Simulated Track Datasets with Experimental Track Statistics from Combo Treatment: Variability in Diffusion Coefficient Between Groups**

(A–D) Joyplots showing distributions of the diffusion parameters α (A), D (B), $D_\alpha$ (C) and $D_{eff}$ (D) together with boxplots of permutation-test p-values obtained from each simulation. Simulations were generated with input diffusion coefficient ($D_i$) of 0.27, 0.31, 0.36, 0.40, and 0.44 μm$^2$/s while keeping the Hurst parameter fixed at 0.35.

Null simulated datasets matched the distributions of track number and track duration obtained during the permutation analysis (Combo; Fig. S3C), whereas the simulated

observation datasets matched the experimental sample size (Combo; Fig. S3B) but did not preserve spatial organization. p-value was 0.05 for power and FPR estimates.

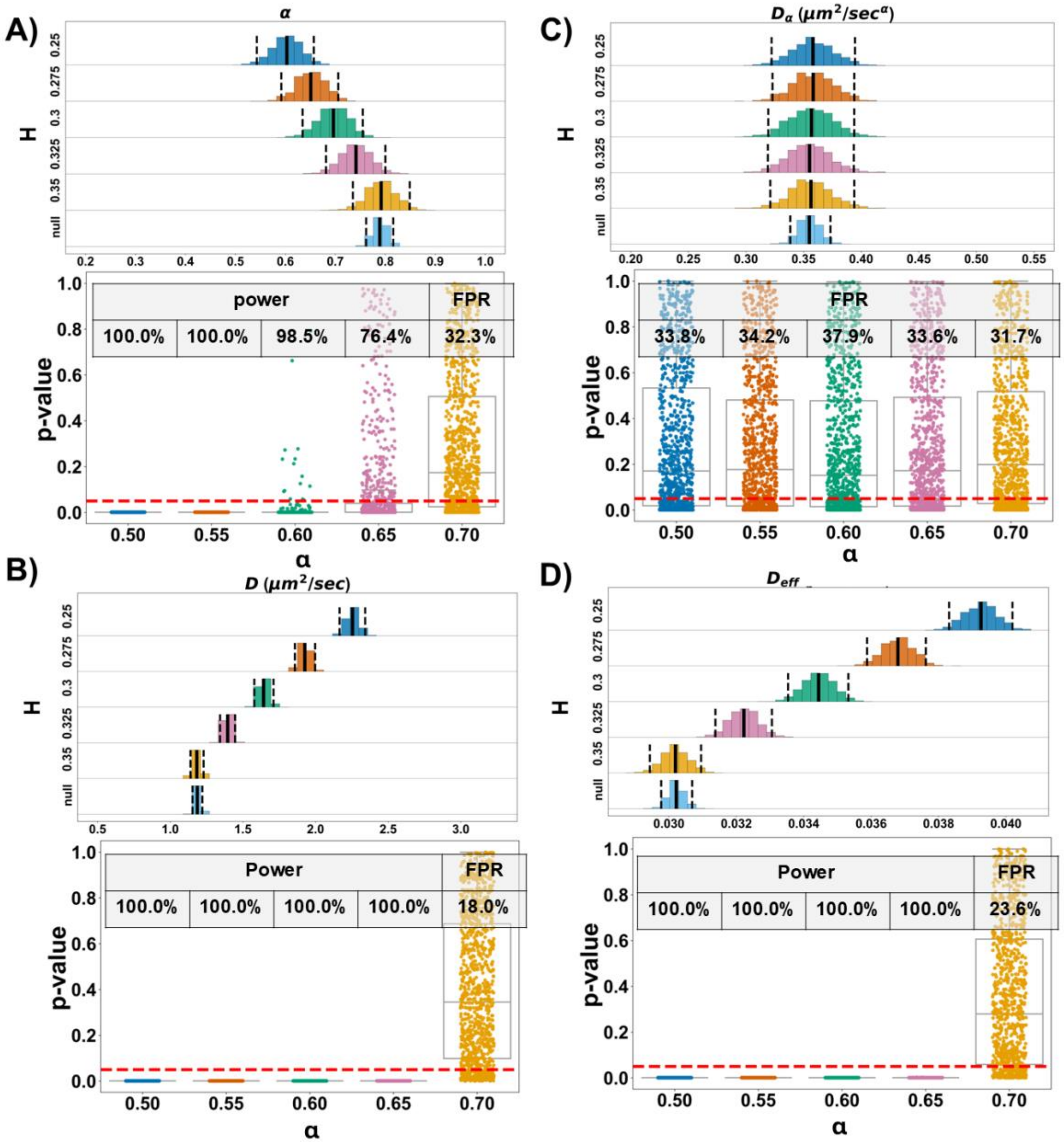


**Figure S12. Validation of the SPaCe-MC Permutation Framework Using Simulated Track Datasets with Experimental Track Statistics from $NaAsO_2$ Treatment: Variability in the Hurst Parameter Between Groups**

(A–D) Joyplots showing distributions of the diffusion parameters α (A), D (B), $D_\alpha$ (C) and $D_{eff}$ (D) together with boxplots of permutation-test p-values obtained from each simulation. Simulations were generated with Hurst parameters of 0.25, 0.30, 0.325, and 0.35 (corresponding to α = 0.50, 0.60, 0.65, and 0.70) while keeping the diffusion coefficient fixed at 0.44 $\mu m^2/s$.

Null simulated datasets matched the distributions of track number and track duration obtained during the permutation analysis (Combo; Fig. S3C), whereas the simulated observation datasets matched the experimental sample size (Combo; Fig. S3B) but did not preserve spatial organization. p-value was 0.05 for power and FPR estimates.

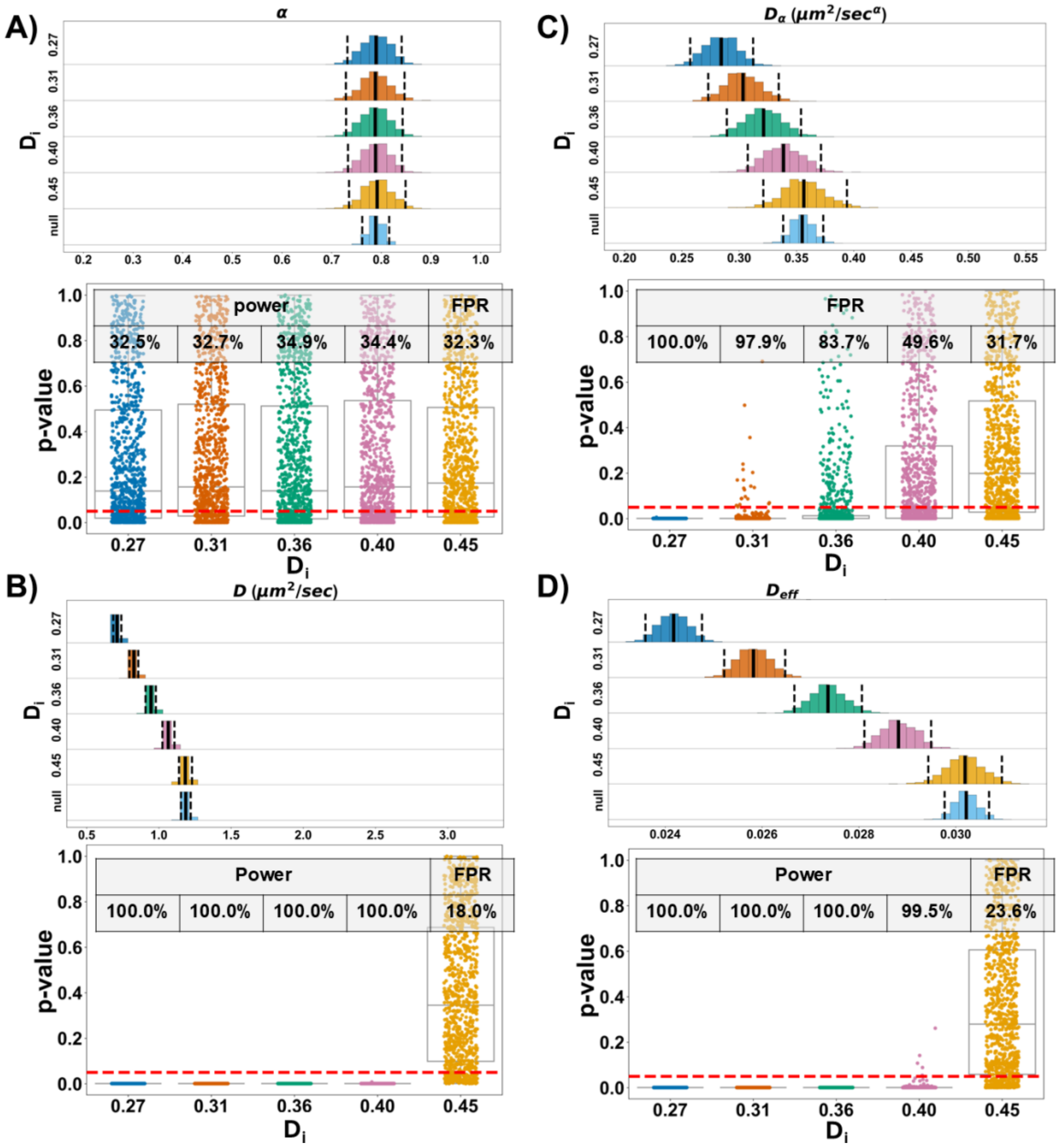


**Figure S13. Validation of the SPaCe-MC Permutation Framework Using Simulated Track Datasets with Experimental Track Statistics from $NaAsO_2$ Treatment: Variability in Diffusion Coefficient Between Groups**

(A–D) Joyplots showing distributions of the diffusion parameters α (A), D (B), $D_\alpha$ (C) and $D_{eff}$ (D) together with boxplots of permutation-test p-values obtained from each simulation. Simulations were generated with input diffusion coefficient ($D_i$) of 0.27, 0.31, 0.36, 0.40, and 0.44 $\mu m^2/s$ while keeping the Hurst parameter fixed at 0.35.

Null simulated datasets matched the distributions of track number and track duration obtained during the permutation analysis (Combo; Fig. S3C), whereas the simulated

observation datasets matched the experimental sample size (Combo; Fig. S3B) but did not preserve spatial organization. p-value was 0.05 for power and FPR estimates.

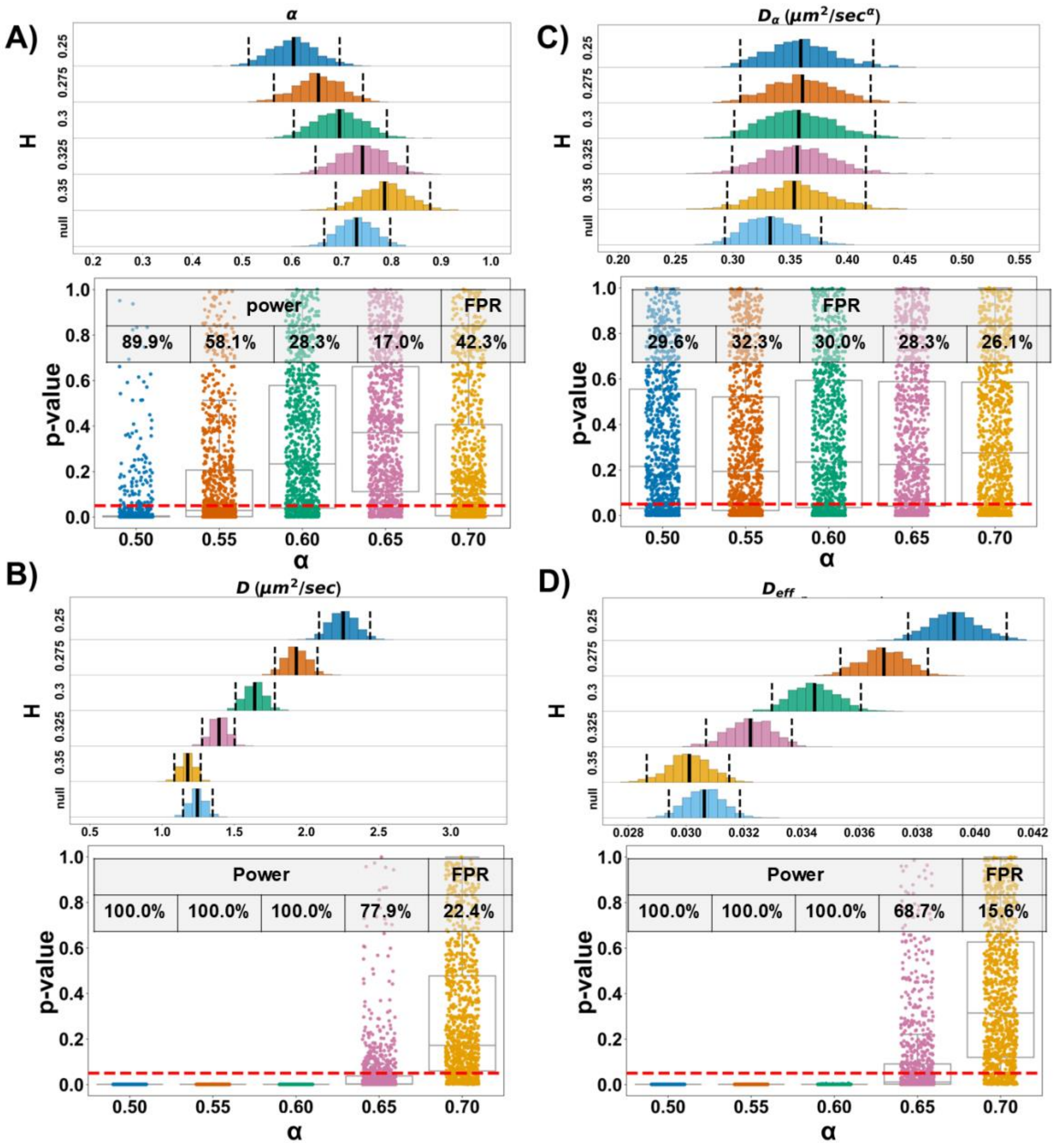


**Figure S14. Validation of the SPaCe-MC Permutation Framework Using Simulated Track Datasets with Experimental Track Statistics from RK-33 Treatment: Variability in the Hurst Parameter Between Groups**

(A–D) Joyplots showing distributions of the diffusion parameters α (A), D (B), $D_\alpha$ (C) and $D_{eff}$ (D) together with boxplots of permutation-test p-values obtained from each simulation. Simulations were generated with Hurst parameters of 0.25, 0.30, 0.325, and 0.35 (corresponding to α = 0.50, 0.60, 0.65, and 0.70) while keeping the diffusion coefficient fixed at 0.44 $\mu m^2/s$.

Null simulated datasets matched the distributions of track number and track duration obtained during the permutation analysis (Combo; Fig. S3C), whereas the simulated observation datasets matched the experimental sample size (Combo; Fig. S3B) but did not preserve spatial organization. p-value was 0.05 for power and FPR estimates.

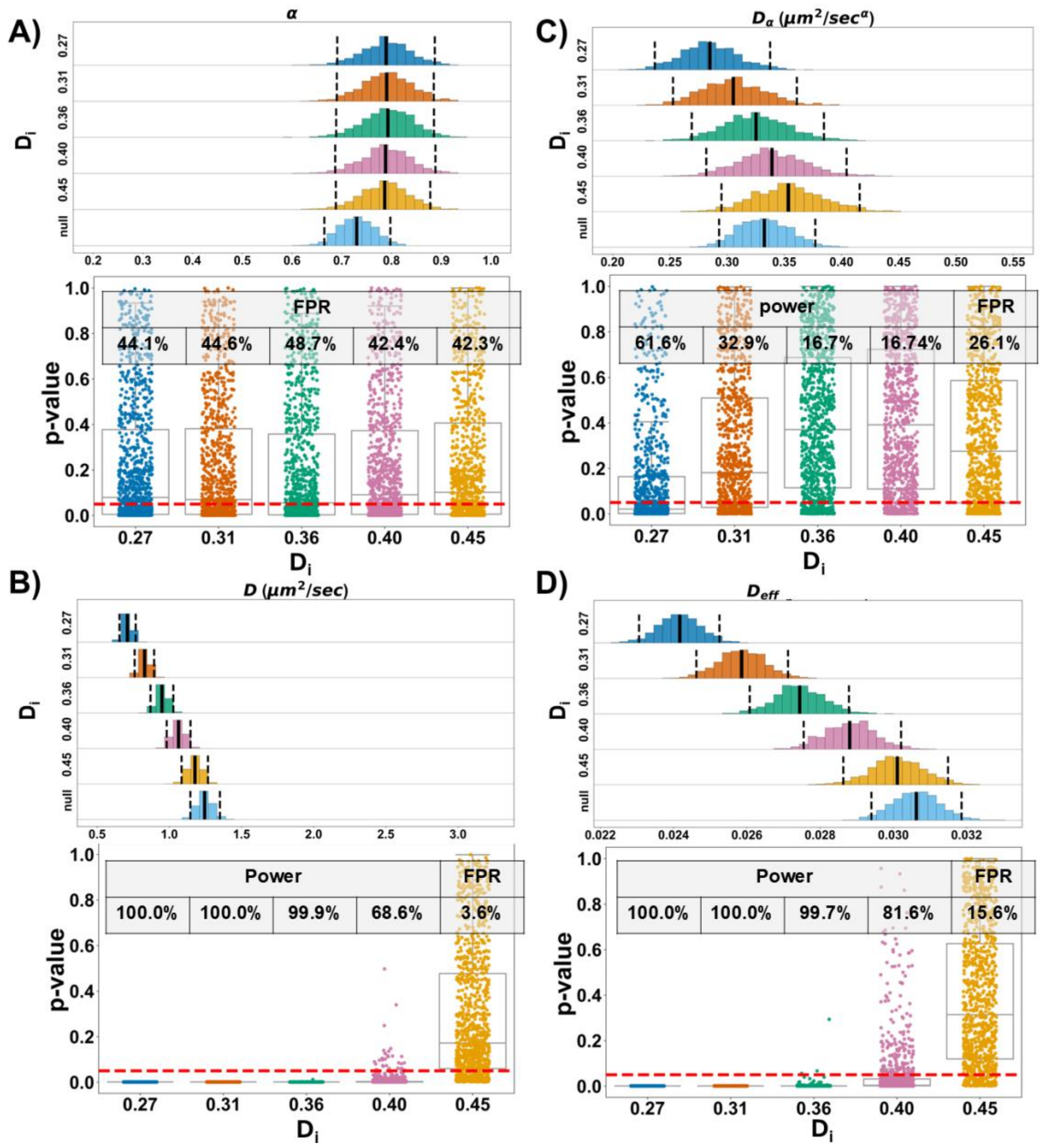


**Figure S15. Validation of the SPaCe-MC Permutation Framework Using Simulated Track Datasets with Experimental Track Statistics from RK-33 Treatment: Variability in Diffusion Coefficient Between Groups**

(A–D) Joyplots showing distributions of the diffusion parameters α (A), D (B), $D_\alpha$ (C) and $D_{eff}$ (D) together with boxplots of permutation-test p-values obtained from each simulation. Simulations were generated with input diffusion coefficient ($D_i$) of 0.27, 0.31, 0.36, 0.40, and 0.44 μm$^2$/s while keeping the Hurst parameter fixed at 0.35.

Null simulated datasets matched the distributions of track number and track duration obtained during the permutation analysis (Combo; Fig. S3C), whereas the simulated

observation datasets matched the experimental sample size (Combo; Fig. S3B) but did not preserve spatial organization. p-value was 0.05 for power and FPR estimates.

A)

| index | Group Number | Hurst parameter (H) | Diffusion Coefficient ($\mu m^2$/sec) | track number | Track duration (ms) |
|---|---|---|---|---|---|
| 0 | 30 | 0.25 | 0.44 | 1000 | 75 |
| 1 | 30 | 0.35 | 0.44 | 1000 | 60 |
| 2 | 30 | 0.35 | 0.22 | 1000 | 75 |
| 3 | 30 | 0.35 | 0.67 | 1000 | 75 |
| 4 | 30 | 0.35 | 0.44 | 1000 | 75 |
| 5 | 30 | 0.35 | [0.22, 0.67] | 1000 | 75 |
| 6 | 30 | 0.35 | 0.44 | 1000 | 90 |
| 7 | 30 | 0.35 | 0.44 | 1000 | [65, 90] |
| 8 | 30 | 0.35 | 0.44 | 2000 | 75 |
| 9 | 30 | 0.35 | 0.44 | 500 | 75 |
| 10 | 30 | 0.35 | 0.44 | [500, 2000] | 75 |
| 11 | 30 | 0.45 | 0.44 | 1000 | 75 |
| 12 | 30 | [0.25, 0.45] | 0.44 | 1000 | 75 |

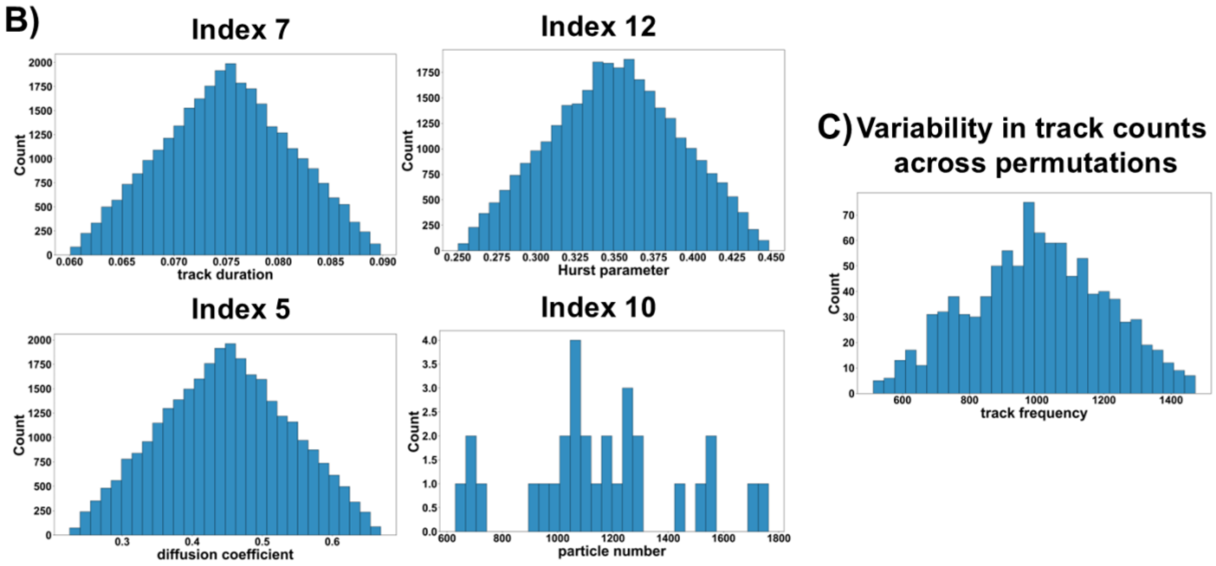


**Figure S16. Description of simulations.**

**(A)** To generate each SPaCe-MC null distribution derived from cytoplasmic tracks, we performed simulations consisting of 30 groups. Within each group, particle trajectories followed a fractional Brownian motion model with either fixed or variable Hurst exponent, diffusion coefficient, track number, and track duration. Variability in the Hurst exponent, diffusion coefficient, and track duration was introduced within groups, whereas variability in track number was introduced between groups. The ranges of track number and track duration were selected to match the experimental distributions shown in Figure S2 C. Approximate diffusion coefficients and Hurst exponents were chosen based on the diffusion parameters estimated from experimental data (Figure 1).

**(B)** Histograms showing the distributions of simulation parameters for simulations with variability: Hurst exponent, diffusion coefficient, track duration, and track number. The x-axis index corresponds to the simulation number.

**(C)** Histograms showing the distribution of track counts across permutations used for "SPaCe-MC-like" analyses in simulations that included variability in track number over permutations.

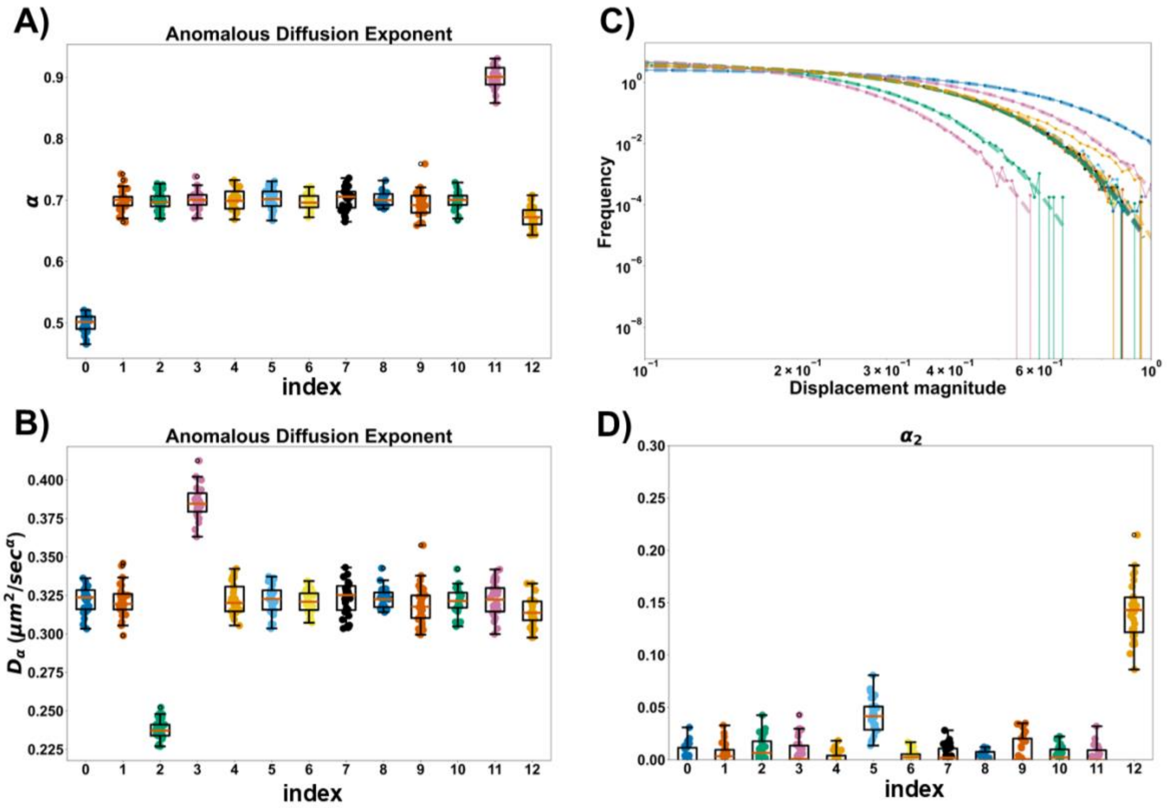


**Figure S17. Estimation of diffusion parameters over simulations**

**(A–B)** Diffusion parameters extracted from anomalous Brownian motion fits: **(A)** anomalous exponent (**α**) and **(B)** anomalous diffusion coefficient (**Dα**). The index number corresponds to the simulation dataset shown in Figure S16.

**(C)** Tails of the GEM particle displacement distributions (10 ms lag time) shown on a log–log scale with Gaussian fits, and **(D)** the corresponding non-Gaussian parameter (**$\alpha_2$**) of the particle displacement distributions.

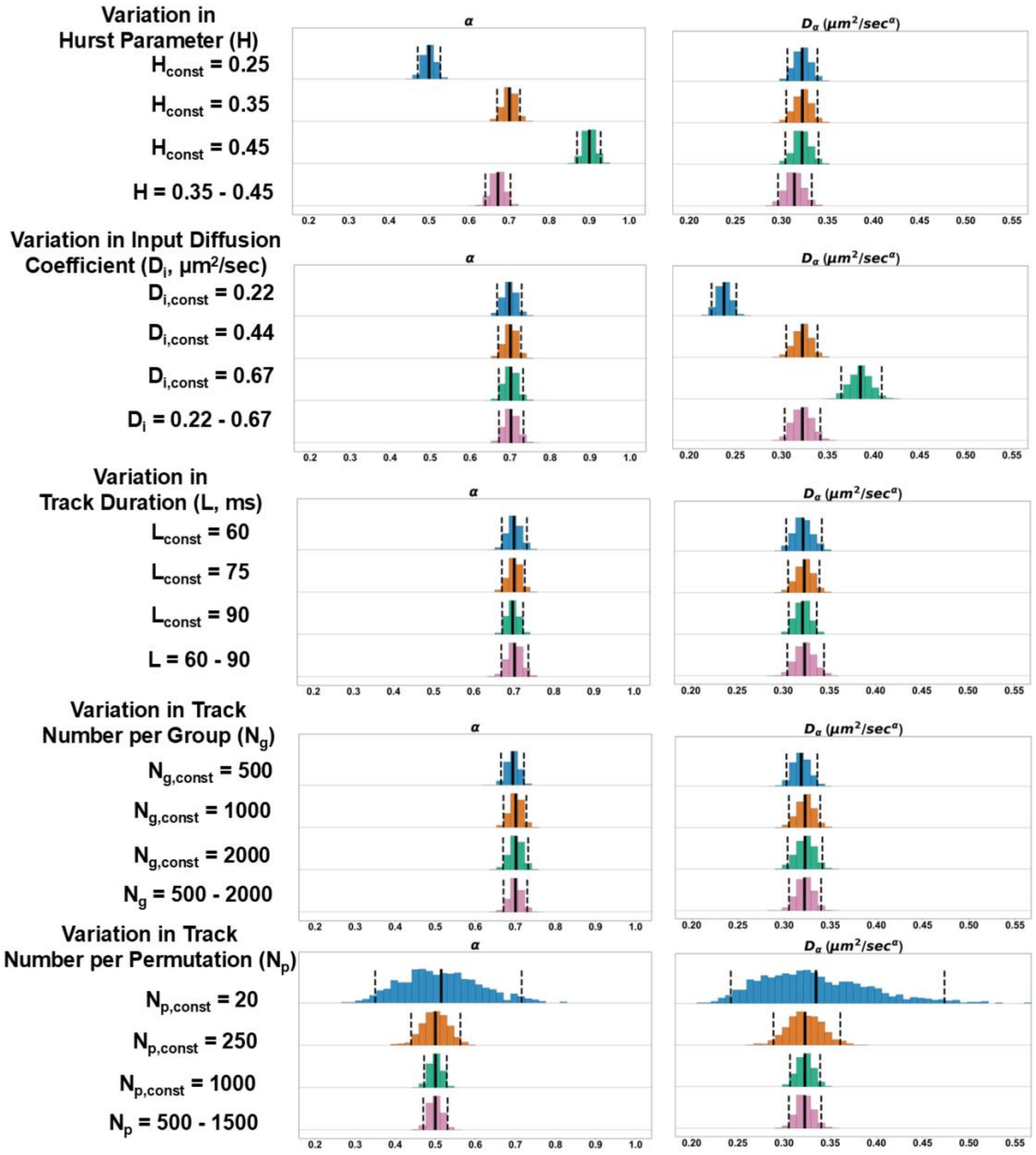


**Figure S18. Effects of simulation parameters on null distributions of diffusion estimates**

Null distributions of diffusion parameters (α and $D_\alpha$) generated using the different statistical properties, including track duration, track number per group, and track number per permutation, and different diffusion parameters, including type of motion and input diffusion coefficient. The row labels indicate the simulation parameter that was varied, whereas the y-axis labels indicate whether the corresponding parameter was fixed or allowed to vary during the simulations.

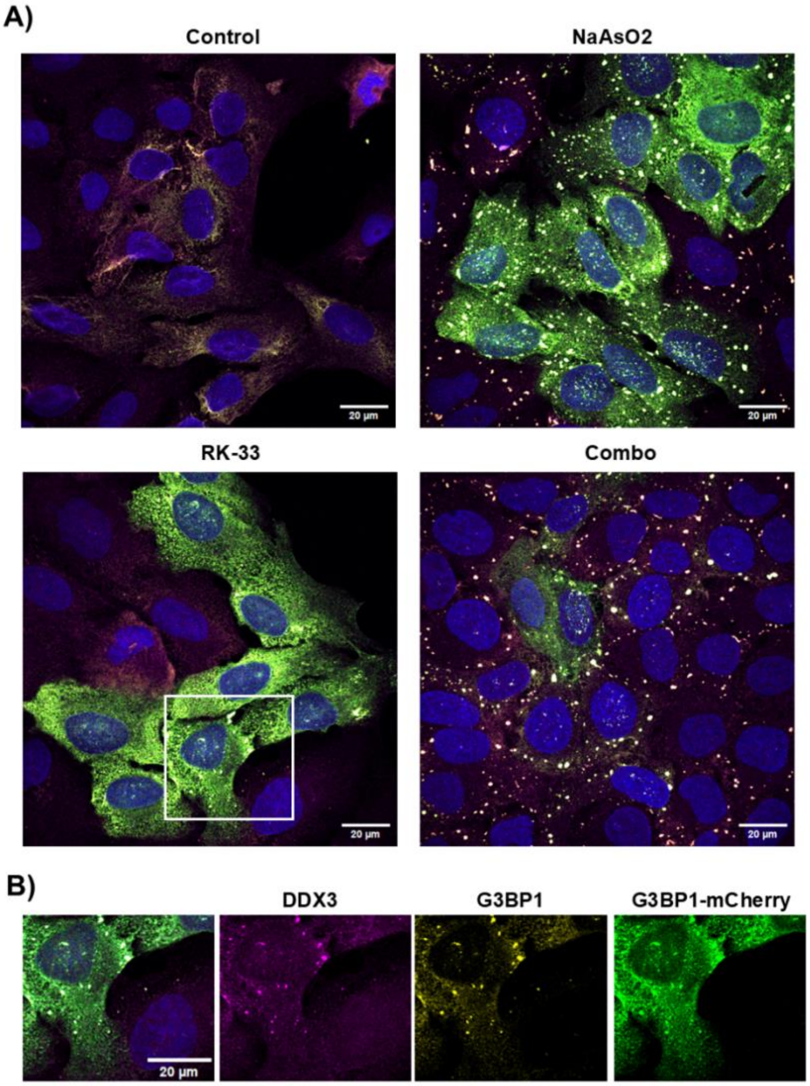


**Figure S19. U2OS cell line expressing G3BP1-mCherry**

**(A)** Maximum-intensity z-projections of U2OS cells and normalized cytofluorograms. DDX3 (magenta), G3BP1-mCherry (green),G3BP1 (yellow), nuclei (blue). Merged images are shown.

**(B)** Cropped single-cell images with merged and individual fluorescence channels. White arrows indicate stress granules positive for both G3BP1 and DDX3.

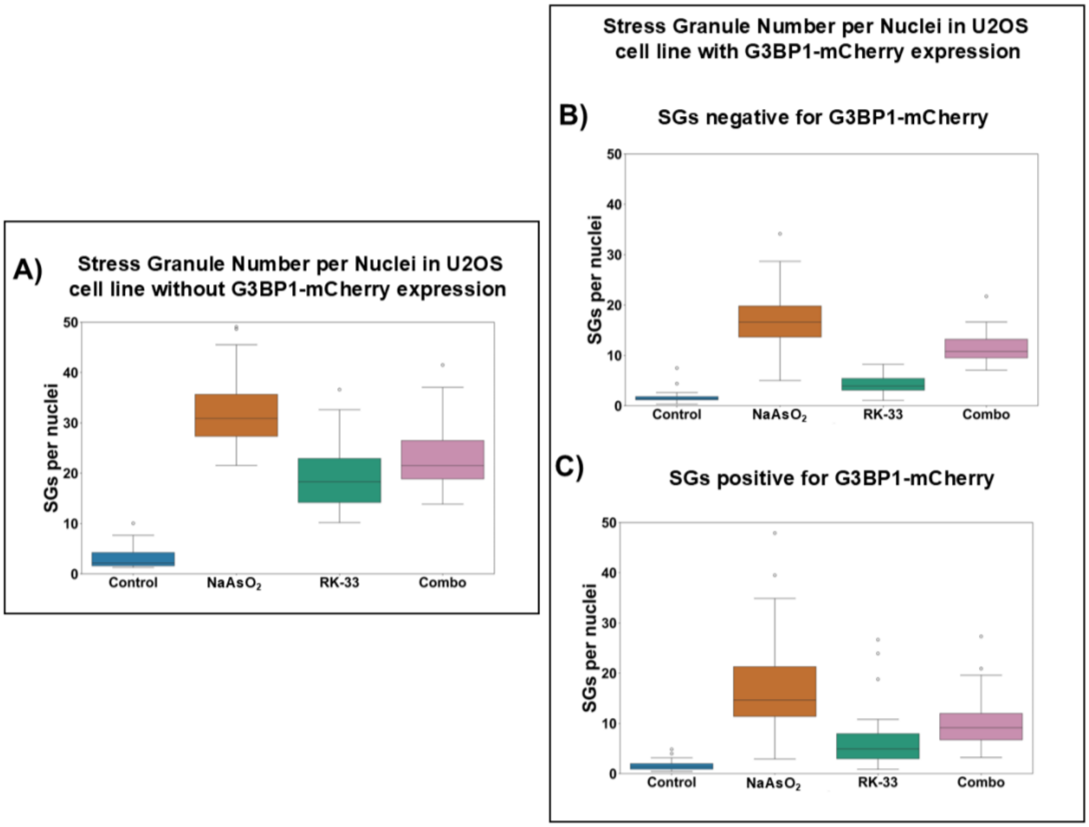


**Figure S20. Quantification of stress granules (SGs) per nucleus in U2OS cells under different G3BP1-mCherry expression conditions.**

Each data point in the boxplot represents the ratio of SG number to nuclei number within a single z-stack. A total of 10 images were analyzed per experimental condition.

**(A)** SGs in U2OS cells with inducible GEM expression lacking G3BP1-mCherry.

**(B)** SGs in U2OS cells with inducible GEM expression and G3BP1-mCherry, including SGs negative for G3BP1-mCherry signal.

**(C)** SGs in U2OS cells with inducible GEM expression and G3BP1-mCherry, including SGs positive for G3BP1-mCherry signal.

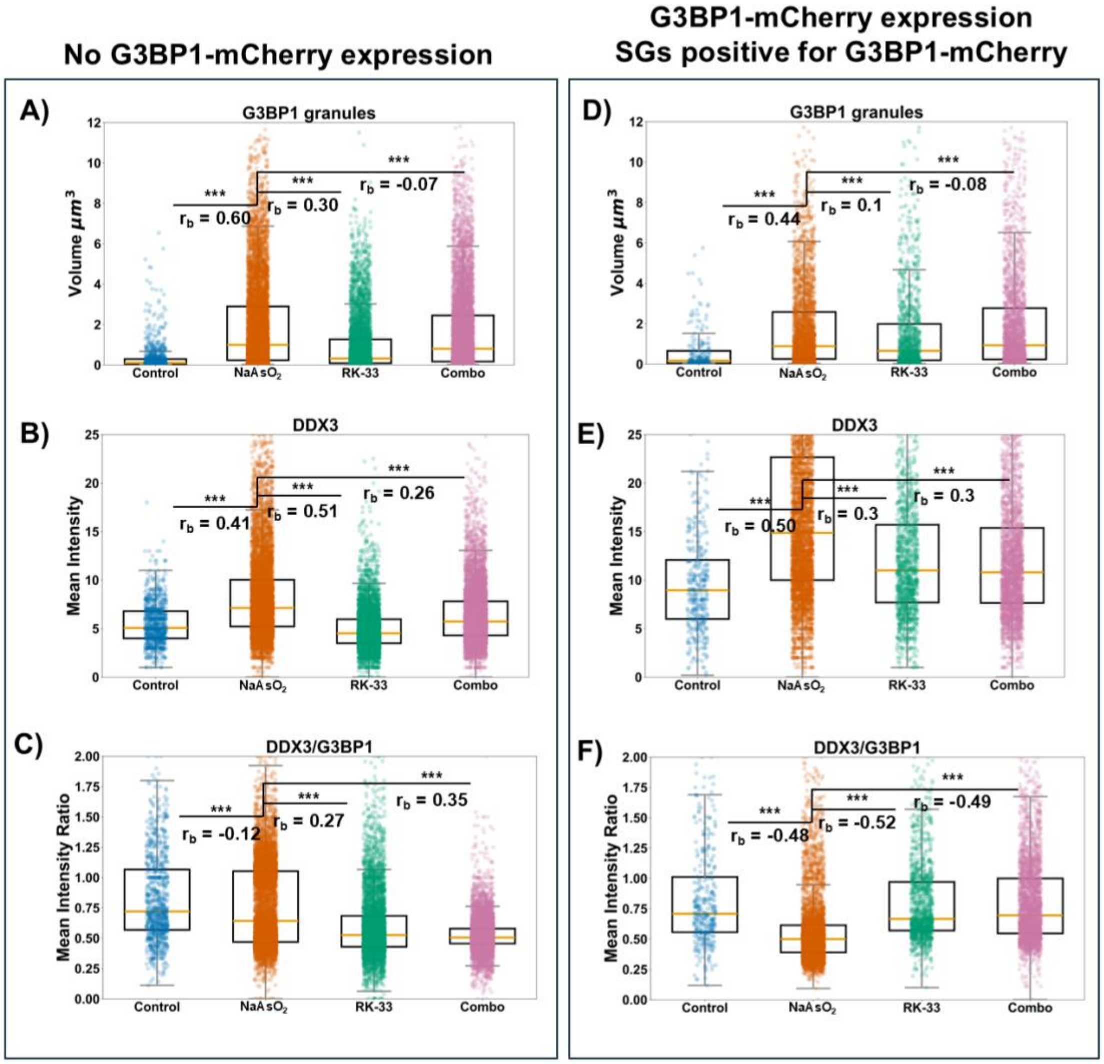


**Figure S21. Quantitative 3D analysis of stress granule formation with DDX3/G3BP1 staining.**

**(A–F)** Quantitative analysis of stress granules (SGs) based on 3D segmentation of z-stack images using the G3BP1 and DDX3 signal. **(A, D)** SG volume distribution. **(B, E)** Mean DDX3 fluorescence intensity per SG. DDX3 intensities were quantified within G3BP1-defined SG boundaries. **(C, F)** Ratio of DDX3 to G3BP1 integrated fluorescence intensity per granule, quantified using 3D z-stack segmentation.

U2OS cell line **(A-C)** without and **(D-F)** with G3BP1-mCherry expression. Each blue dot represents a single granule. The Mann-Whitney U test ($*p < 0.05$, $**p < 0.01$, $***p < 0.001$). Effect size was measured as rank biserial rb: : rb < 0.3 small, 0.3 < rb < 0.5 medium, rb > 0.5 large.
